# Abscisic acid signaling gates salt-specific responses of plant roots

**DOI:** 10.1101/2023.12.28.572987

**Authors:** Jasper Lamers, Yanxia Zhang, Eva van Zelm, A. Jessica Meyer, Thijs de Zeeuw, Francel Verstappen, Mark Veen, Ayodeji O. Deolu-Ajayi, Charlotte M.M. Gommers, Christa Testerink

## Abstract

Soil salinity presents a dual challenge for plants, involving both osmotic and ionic stress. In response, plants deploy distinct yet interconnected mechanisms to cope with these facets of salinity stress. In this investigation, we observed a substantial overlap in the salt (NaCl)-induced transcriptional responses of Arabidopsis roots with those triggered by osmotic stress or the plant stress hormone abscisic acid (ABA), as anticipated. Notably, a specific cluster of genes responded uniquely to sodium (Na^+^) ions. Surprisingly, expression of sodium-induced genes exhibited a negative correlation with the ABA response and preceded the activation of genes induced by the osmotic stress component of salt. Elevated exogenous ABA levels resulted in the complete abolition of sodium-induced responses. Consistently, ABA signalling mutants displayed prolonged sodium-induced gene expression, coupled with increased root cell damage under high salinity conditions. Moreover, ABA signalling mutants were unable to redirect root growth to avoid high sodium concentrations and failed to contain their root cell swelling in the presence of elevated salt levels.

In summary, our findings unveil an unexpected and pivotal role for ABA signaling in mitigating cellular damage induced by salinity stress and modulating sodium-specific responses in plant roots.

## Introduction

Plants are bound to their location and must adequately respond to their environment to survive. This requires the translation of environmental cues to cellular signaling, which is achieved by various sensing mechanisms. Soil salinization is one of the major threats to agriculture affecting >1 billion hectares worldwide, including 20% of all irrigated farmlands^1^.

High soil salinity lowers the water potential, which limits plant water content and causes osmotic stress, which is putatively sensed by mechanosensitive Ca^2+^ channels^2^. In addition, toxic ions (sodium and chloride) cause ionic stress due to their accumulation in plants, inhibiting several cellular processes^3,4^. Some plants, including mangrove trees^5^ and quinoa^6^, have evolved mechanisms to deal with extremely high salt concentrations, and are called halophyte species. On the other hand, the majority of plants, including most important crops, are highly salt sensitive (termed glycophyte species)^3^. Interestingly, glycophytes can redirect root growth to specifically avoid high sodium concentrations in soil^7^, a process named halotropism, which is not induced by an equimolar level of osmotic stress or by other monovalent cations, such as K^+^ or Li^+^ ^7–9^. This indicates the existence of a yet unexplored sodium-induced sensing mechanism in plants that allows plants to mount responses to this toxic ion that is increasingly present in natural and agricultural soils^1^. On the other hand, many responses to salt overlap with those induced by osmotic stresses, including drought. The plant hormone abscisic acid (ABA) is produced in response to many water-limiting abiotic stresses (such as drought, osmotic stress, salinity, cold and frost) and has an important role in the regulation of germination, stomatal closure^10^ and root development^11^. ABA is sensed by binding to the *PYRABACTIN RESISTANCE1 (PYR)/PYR1-LIKE* family of receptors^12^, which releases the inhibition by PROTEIN PHOSPHATASES TYPE 2C (PP2C) proteins^13^ of the SUCROSE NON-FERMENTING 1-RELATED PROTEIN KINASE2 family (SnRK2; 2.2, 2.3 and 2.6 in Arabidopsis^14^; Fig. S1). SnRK2s activate downstream transcription factors (TFs) such as ABA INSENSITIVE 3 (ABI3), ABI4, ABI5 and ABA-RESPONSIVE ELEMENT-BINDING FACTOR/ABSCISIC ACID RESPONSIVE ELEMENT-BINDING FACTOR 1 TFs (AREB/ABF; 4 homologs in Arabidopsis)^15,16^. In response to salt, ABA accumulation and ABA-induced gene expression peak at 3-6 hours after salt application in Arabidopsis roots and decrease again after 24 hours^17,18^, which is similar to the response of plants to water deficit^18^.

Until recently it was widely accepted that the osmotic responses induced by salt precede those induced specifically by the ionic stress, which would occur only after sodium and chloride ions have accumulated to toxic levels^3^. The discovery of the sodium-specific halotropism response^7,9^, and the rapid (<20 seconds) induction of Ca^2+^ waves migrating from root to shoot after root exposure to salt, but not osmotic stress^19,20^, have challenged this longstanding idea.

To understand how plant roots integrate the osmotic and ionic component in the early response to salt, we first sought to identify specific sodium-induced transcriptional responses in Arabidopsis. We performed an RNA sequencing experiment on roots of 8-day-old seedlings with two sodium treatments (NaCl and NaNO_3_), an ionic control (KCl) and an osmotic control (sorbitol) at 6 and 24 hours. We discovered that genes that are specifically induced by sodium ions, are induced earlier than those general to all osmotic stresses and are negatively regulated by ABA. Further investigation of the relation between ABA and sodium-induced responses revealed that endogenous ABA signaling not only plays a general role in abiotic stress responses but is also required to specifically attenuate rapid sodium-induced responses in roots. Timing of ABA accumulation acts to limit sodium-induced cell damage and finetunes salt-induced changes in root growth and morphology.

## Results

### Salinity triggers sodium-induced gene expression that is inhibited by ABA

To identify sodium-induced transcriptional changes, 8-day old Arabidopsis seedlings were subjected to mock (control), 130mM ionic (NaCl, NaNO_3_, KCl), 260mM osmotic (sorbitol) or 25 µM ABA treatments on solid ½MS medium. Roots were harvested after 6 and 24 hours. The principal component analyses (PCA) of both timepoints show that the treatments caused more distinct expression patterns after 6 hours compared to 24 hours (Fig. S2A and B). Next, the ionic (NaCl, NaNO_3_, KCl) and osmotic (sorbitol) treatments were used to identify sodium-induced gene expression. Genes that were differentially expressed (False Discovery Rate (FDR) < 0.05, Log fold change (LFC) > |0.5|) in at least one of the treatments compared to the control were selected for further analysis. The overlap between treatments is visualized in Venn diagrams (Fig. 1A, Fig. S3A). Up- and downregulation were presented separately, and we defined sodium-induced gene regulation as agonistically regulated by both NaCl and NaNO_3_, and not significantly, or antagonistically by KCl and sorbitol treatments.

**Figure 1.**
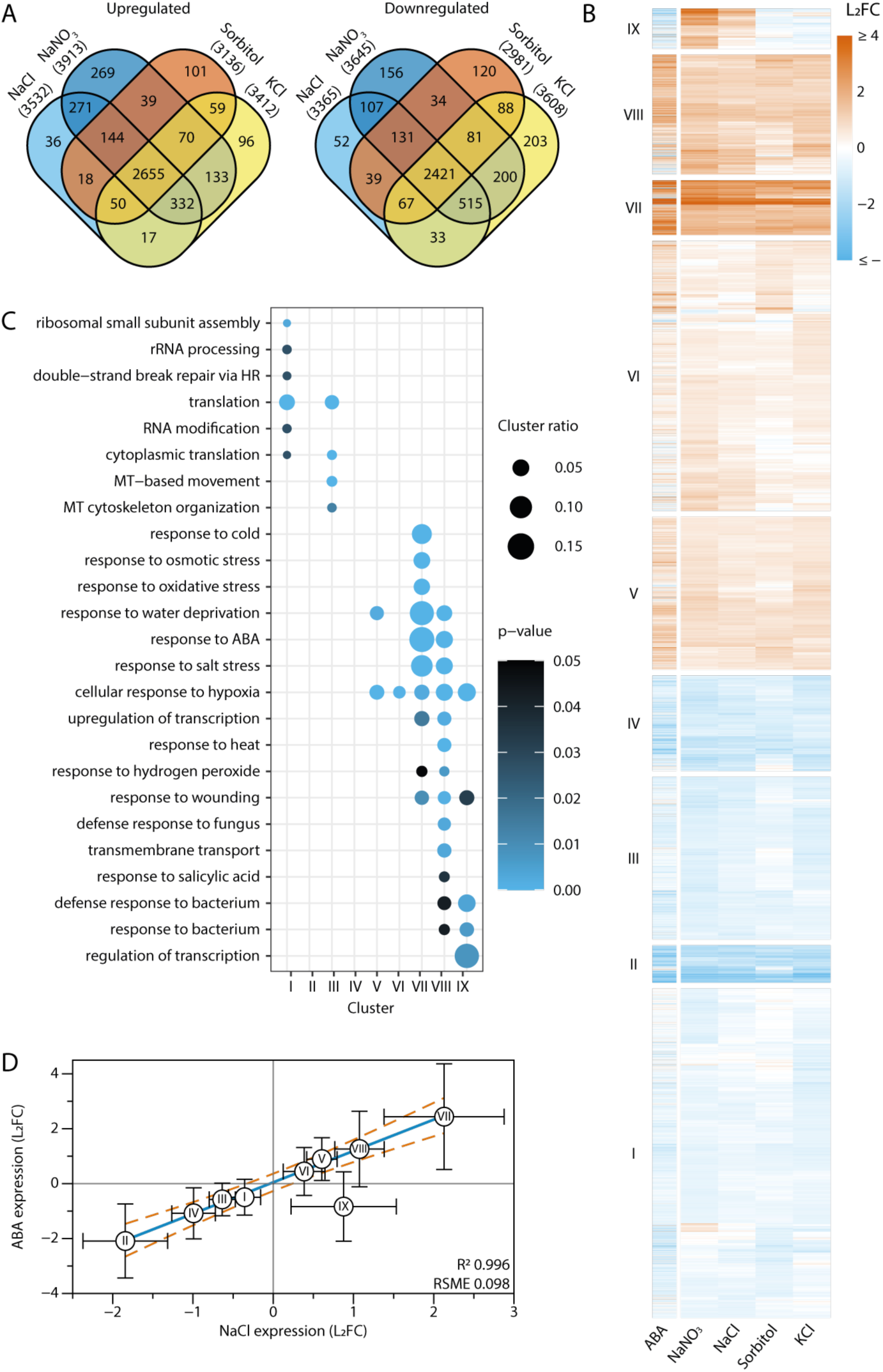
The Arabidopsis root transcriptome response to sodium ions negatively correlates with the response to ABA. Transcriptomics on 8-day-old Arabidopsis roots treated with 130mM NaCl, 130mM NaNO3, 130mM KCl, 260mM Sorbitol or 25µM ABA for 6 hours. Transcripts were filtered for significance (FDR < 0.05) and expression (LFC > |0.5|) and were included when these conditions were met in at least one of the 4 treatments (NaCl, NaNO3, KCl or sorbitol). This resulted in a selection of 8431 DEGs. (A) Venn diagrams of up- and downregulated genes showing the overlap of differential gene expression between the different treatments at 6 hours. (B) Heatmap of the 8431 DEGs. Treatments (columns) and transcripts (rows) were clustered with Euclidean distance mapping and Ward clustering. ABA treatment was not used for clustering and was added as an annotation column. The heatmap was divided into 9 clusters, which is indicated on the left side of the cluster. (C) GOterm enrichment per cluster was analyzed with GOseq^61^. GO terms containing less than 10 genes in total were excluded from the plot. The color indicates the Benjamin Hochberg adjusted p-value. Size indicates the ratio of DEGs in the GO term to the total number of genes in the term. (D) Scatterplot of the mean L2FC in ABA and NaCl per cluster. The linear fit of cluster I-VIII is shown in blue, with a 99.99% prediction plot shown in red lines.

We identified 271 upregulated and 107 downregulated sodium-induced genes at 6 hours (Fig. 1A) and 181 upregulated genes and 75 downregulated genes at 24 hours (Fig. S3A). However, few genes exceeded LFC > |1| in the NaCl treatment at 24 hours (19) compared to 6 hours (80) (Fig. S3B). Hence, we studied the 6-hour dataset in more detail.

The 8431 selected DEGs at 6 hours after treatment were clustered to identify common expression patterns in response to the different treatments. Clustering was performed using Euclidean distance mapping and Ward clustering (Fig. 1B). The ABA treatment was not included in the selection of DEGs or the clustering itself but was added as a separate annotation column. At 6 hours, the genes were divided into 9 clusters (Fig. 1B). Four clusters (I-IV) are predominantly downregulated by all treatments, while four others (V-VIII) are upregulated. On the contrary, on the cluster level, genes in cluster IX show upregulation by sodium (NaCl, NaNO3), but not by sorbitol, KCl or ABA. 125 genes in this cluster were sodium-induced on the individual gene level as identified in the Venn diagrams of Fig. 1A. A gene ontology (GO) term enrichment analysis showed that genes in the downregulated clusters (I-IV) are mostly related to translation, and those in the upregulated clusters (IV-VIII) enriched for terms related to abiotic stresses (Fig. 1C). Surprisingly, GO terms associated with (defense) response to bacterium and wounding, were strongly enriched in the sodium-induced cluster (IX) at 6 hours. Consistent with the PCA plots and the Venn diagrams, no sodium-induced cluster was identified at 24 hours (Fig. S3C), which indicates that a distinct sodium-induced response is transient. The mean NaCl response of genes in cluster I – VIII positively correlates with the response to ABA (R^2^ = 0.996, Fig. 1D), as expected. Interestingly, this correlation is not observed in the mean transcriptional response of cluster IX genes (Residual Standard Deviation > 2.5σ). The genes in this cluster are generally repressed by the ABA treatment. Based on the similarities of clustering results with ABA-induced expression patterns, we hypothesize that an increase in ABA, caused by the osmotic component of NaCl stress, is responsible for the transcriptional changes of genes in cluster I-VIII, but could potentially at the same time compromise the sodium-induced response to salinity (cluster IX).

### Sodium-induced gene expression peaks at early timepoints

We hypothesized that the activation of sodium-induced gene expression and the induction of ABA signaling by high salinity could be temporarily separated. Our transcriptome dataset Cluster IX transcripts generally showed very low abundance in the NaCl treatment compared to other genes and were mostly undetectable in control conditions (Fig. S3D). We first selected the most upregulated sodium-induced gene (*REDOX RESPONSIVE TRANSCRIPTION FACTOR 1; RRTF1/ERF109*) as a robust sodium-induced marker gene and quantified the expression *RRTF1,* and an ABA marker gene *(RESPONSIVE TO ABA 18; RAB18)*^14^ over time after NaCl stress induction by RT-qPCR. Expression of *RRTF1* was induced within an hour after salt application, was quickly reduced at 3 hours, followed by a stable level of expression and a further decrease at 24 hours (Fig. 2A). *RAB18* expression increased gradually upon NaCl treatment and reached its maximum expression level at 3 hours post-salt application and stayed elevated for several hours (Fig. 2B). To investigate how these expression patterns relate to the synthesis of ABA in roots, we quantified ABA levels at 1, 3, 6 and 12 hours after NaCl, KCl and sorbitol treatment. ABA levels significantly increased after 1 hour, peaked at 3 hours for all three stress treatments and slowly declined afterwards (Fig. 2C). Taken together, these results indicate that the early sodium-induced gene expression is most apparent 1 hour after stress, and is later reduced, while ABA levels are still increasing.

**Figure 2.**
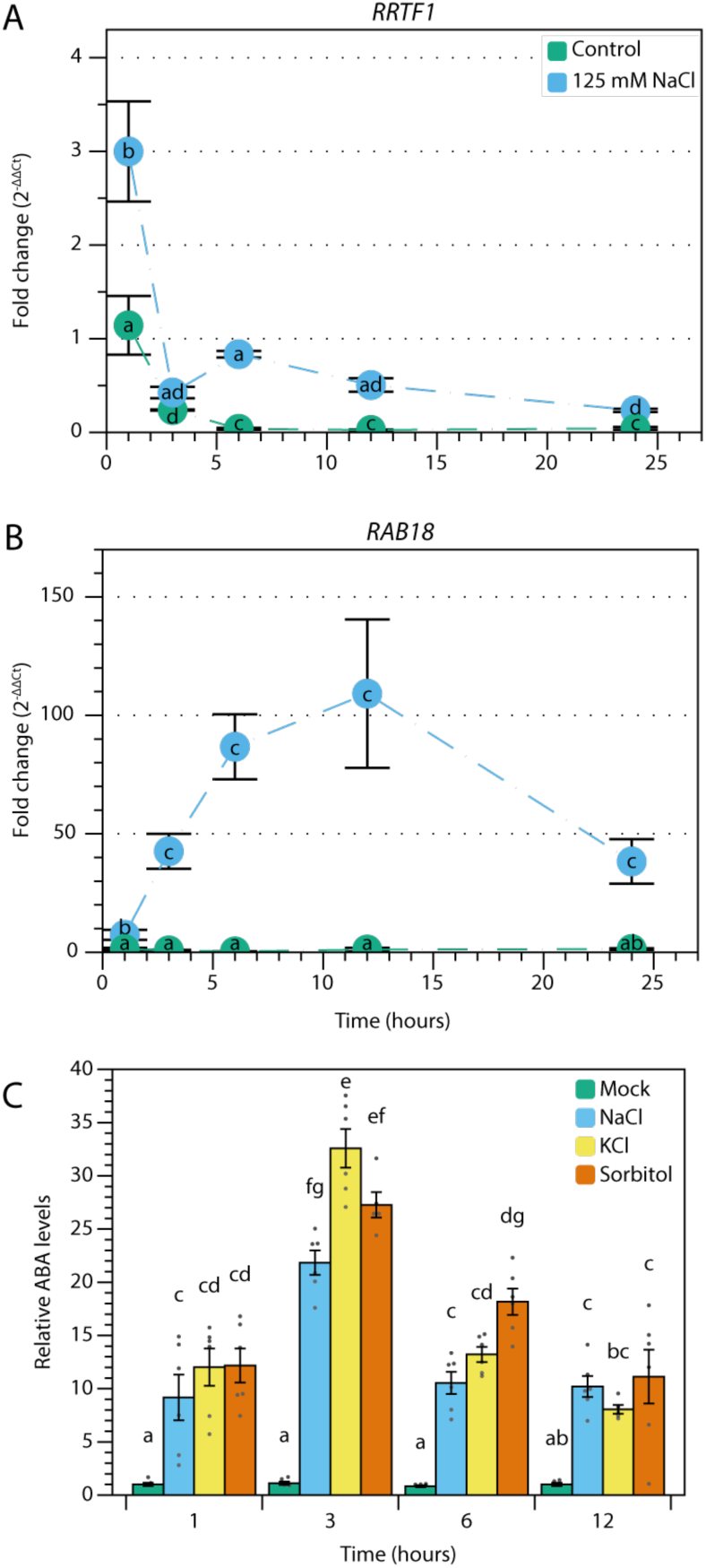
Sodium-induced gene expression precedes ABA responses. (A-B) Relative gene expression of a sodium-induced gene (*RRTF1*, A) and an ABA signaling marker (*RAB18*, B) over time in roots of 7-day-old seedlings after transfer to control or 125mM NaCl medium. Expression of all samples was normalized to the 1-hour control condition using *MON1* as a housekeeping gene. Dots represent the mean (n=4) +/-SE. Data was statistically analyzed using two-way ANOVA, followed by a Tukey post-hoc test. Different letters indicate significant differences (p < 0.05). (C) Normalized ABA levels compared to the mock treatment at 1 hour over time in roots of 7-day-old seedlings after transfer to 125mM NaCl, 125mM KCl, or 250mM sorbitol. Bars represent mean values (n = 5-6) +/-SE and individual datapoints are shown as dots. Data was statistically analyzed using two-way ANOVA, followed by a Tukey post-hoc test. Different letters indicate significant differences (p < 0.05).

### ABA pretreatment inhibits sodium-induced responses

We have seen that (1) sodium-induced genes are repressed by ABA in the absence of NaCl and (2) NaCl-induced ABA signaling peaks at later timepoints. Hence, we questioned if ABA can also suppress sodium-induced gene expression in the presence of NaCl. To test this, we applied a combined treatment of ABA and NaCl. For these experiments, we selected a lower concentration of 5 µM ABA, which approximates gene expression changes of ABA marker genes as did 125 mM NaCl (Fig. S4). To maximize ABA-induced gene expression at the start of the NaCl application (Fig. 2B), seedlings were transferred to mock or 5µM ABA treatment plates 12 hours prior to the NaCl or control treatment with the corresponding ABA concentration. Gene expression was quantified for *RRTF1* and the ABA signaling marker gene *RAB18*. *RRTF1* was significantly (p < 0.05) upregulated after 1 hour of NaCl, while a pretreatment with 5 µM ABA significantly reduced NaCl-induced expression of *RRTF1* to control levels (Fig. 3A). As expected, the

**Figure 3.**
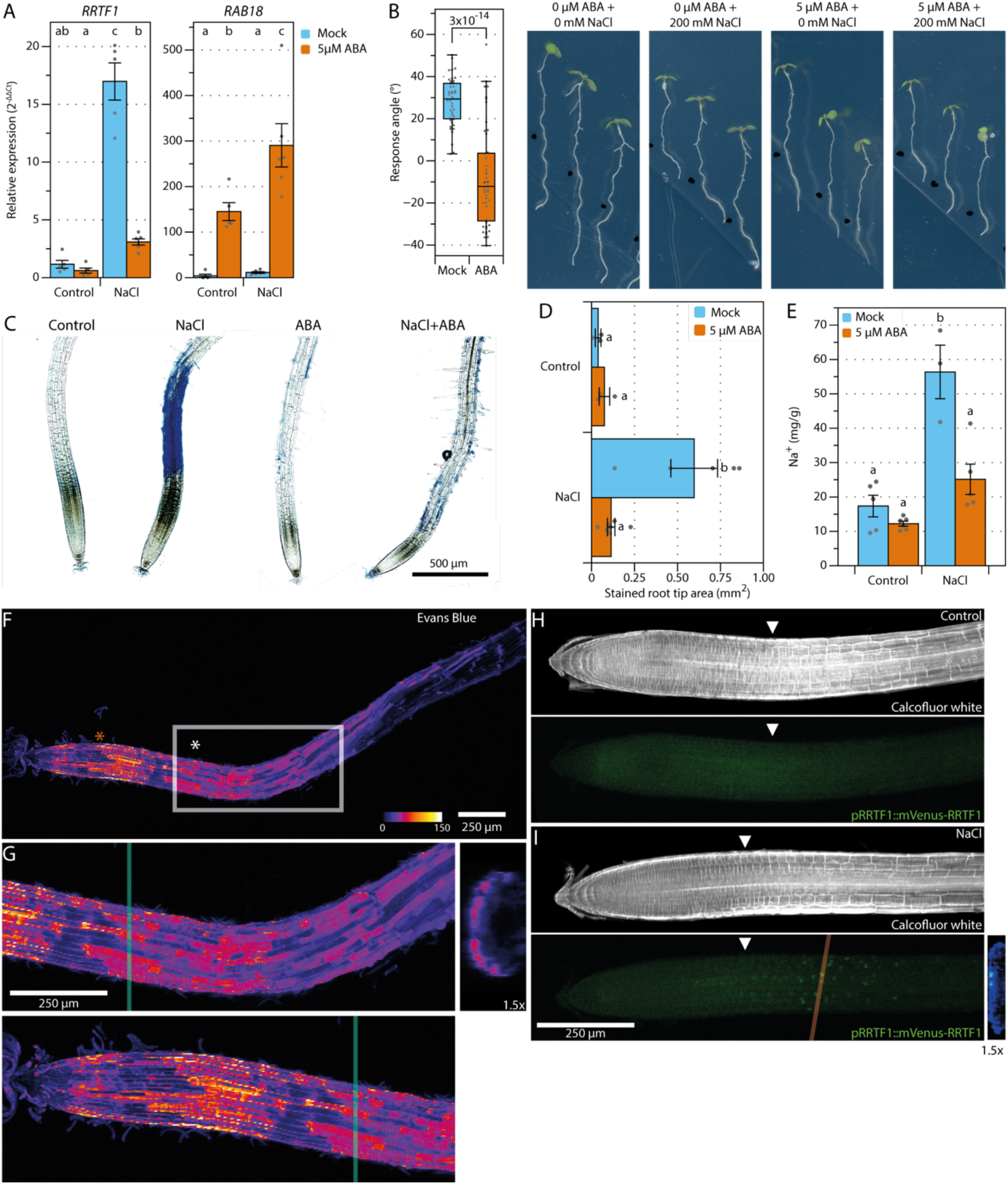
ABA represses sodium-induced responses. (A) Relative gene expression of 2 sodium-induced transcripts (*RRTF1*, *CBF1*) and an ABA signaling marker (*RAB18*) in roots of 7-day-old seedlings after transfer to control or 125mM NaCl medium for 1 hour and with (orange) or without (blue) 5µM ABA pretreatment for 12 hours. Expression was normalized to the control condition without ABA. Bars represent mean values (n = 5-6) +/-SE and individual datapoints are shown as dots. Statistical analysis was performed using non-parametrical Dunn’s test on the log2 transformed data and corrected for multiple testing using Benjamin-Hochberg (BH) procedures. Letters indicate statistical groups. (B) Quantification and representative images of the halotropism response at 48 hours after introduction of the gradient in the presence or absence of an ABA pretreatment. The response angle shows the difference between control and NaCl conditions for the respective ABA treatment. The relative growth shows the ratio between NaCl and control conditions for the respective ABA. Statistics were performed using pairwise t-tests. (C) Representative images of Evans Blue staining for plasma membrane damage at 48 hours after the introduction of the gradient of the halotropism assay. (D) The affected area of the assay (C) was quantified using a script (Fig. S15). Bars represent mean values (n = 5) +/-SE and individual datapoints are shown as dots. Different letters indicate significant differences, analyzed by one-way ANOVA, followed by a Tukey post-hoc test, p < 0.05. (E) Sodium contents in roots of 7-day-old seedlings after transfer to control or 125mM NaCl medium for 6 hours and with (red) or without (blue) 5µM ABA pretreatment for 12 hours. Bars represent mean values (n = 3-6) +/-SE and individual datapoints are shown as dots. (F-G) Confocal images of an Evans Blue stained root at 24 hours after 125mM NaCl treatment. The red asterisk indicates superficial staining of Evans Blue, which is also observed in control conditions (see Figure S6A/B). The white asterisk indicates sodium-induced cell damage that penetrates to deeper cell layers and is absent from the control samples. (G) Enlargement of Fig. 3F (marked with the white box). The orthogonal view (on the green line) is 3x enlarged and shown on the side. (H-I) Maximum projection confocal images of roots expressing *pRRTF1::RRTF1-Venus* at 2 hours after control (H) or 125mM NaCl treatment (I). The orthogonal view (on the red line) is 1.5x enlarged and shown on the side. The midplane (1 stack) of the calcofluor-white cell wall staining is shown in white. White arrowheads indicate the start of the elongation zone based on the cortical cell length.

ABA signaling marker gene *RAB18* was not induced after 1 hour of salt stress, but upregulated by ABA and even more upregulated by the combination of ABA and NaCl. Taken together, these results suggest that the strong induction of the sodium-induced marker gene *RRTF1* at 1 hour after salt treatment is facilitated by the low ABA levels at this stage (Fig. 2C).

To investigate whether ABA also represses sodium-induced phenotypic responses of roots, we next studied halotropism, which is the redirection of root growth to specifically avoid high sodium concentrations^7,9^. Four-day old seedlings were transferred to fresh plates with or without 5 µM ABA and 12 hours later a salt gradient was introduced by removing the lower part of the agar and replacing it with control medium or medium supplemented with 200mM NaCl (plus 5 µM ABA for ABA pretreated samples). After 24 and 48 hours, the NaCl gradient caused a change in root growth direction of almost 30°, consistent with published data^9^ (Fig. 3B/S5). Intriguingly, ABA pretreated roots were initially avoiding high sodium concentrations, but showed positive halotropism, bending towards the higher NaCl concentration after 48 hours (Fig. 3B). Because the sodium-induced gene cluster IX is enriched for genes involved in wounding-related processes (Fig. 1D), we investigated whether salinity stress causes wounding to roots. Cell damage of root cells was assessed using Evans Blue staining^21^ at 48 hours after a halotropism assay in the presence or absence of an ABA pretreatment. No damage was detected in the control group, but the NaCl-treated roots were severely damaged in the elongation zone (Fig. 3C) and ABA reduced the NaCl-induced damage (Fig. 3C/D).

As the ABA pretreatment abolishes sodium-induced responses, we hypothesized that ABA lowers Na^+^ levels. Therefore, we quantified sodium content in 7-day-old roots at 6 hours after a homogeneous control or 125mM NaCl treatment, in the presence or absence of a 5 µM ABA pretreatment. ABA pretreatment strongly reduced the sodium content in the root (Fig. 3E), which might cause the inhibition of sodium-induced root growth responses.

To visualize NaCl-induced cell damage at a cellular resolution, we imaged NaCl-treated roots with Evans Blue using confocal microscopy. We confirmed staining of cells in the early elongation zone in response to salt treatment (Fig. 3F/G and Fig. 6F). As an ABA pretreatment abolishes both NaCl-induced cell damage and sodium-induced gene expression, we investigated if both processes are spatially overlapping. Therefore, we visualized the localization of *pRRTF1:RRTF1-Venus*^22^ at 2 hours after salt stress and compared it to the confocal images of the Evans Blue staining. In accordance, RRTF1-Venus accumulated in the epidermal cells of the early elongation zone (600-850 µm from the root tip) in response to salt stress (Fig, 3H/I). Thus, only few cells express the sodium-induced *RRTF1* marker, which aligns well with the low transcript counts in the RNA sequencing dataset as described above (Fig. 1/S3). In conclusion, ABA consistently abolishes sodium-induced responses, including gene expression, halotropism and cell damage of the elongation zone of roots exposed to high salinity. As salt stress induces cell damage, we investigated the role of Jasmonic Acid (JA), a plant hormone involved in plant wounding responses, in sodium-induced responses. JA is also known to quickly induce *RRTF1* expression during mechanical damage^23,24^. Even though control levels of *RRTF1* were significantly

lower in mutants with reduced biologically active JA (*aos* and *jar1-1*), expression was still induced by NaCl (Fig. 4A). These mutants also showed wildtype root halotropic reponses (Fig. 4A/B/S7)., indicating that early sodium-induced responses are JA-independent. As reactive oxygen species (ROS) are quickly induced by wounding^25^ and *RRTF1* expression is induced by ROS^26^, we hypothesized that ROS could be the signal responsible for the induction of sodium-induced gene expression. NADPH RESPIRATORY BURST OXIDASE PROTEINs (RBOHs) are a major source for ROS production during NaCl stress^27^. We investigated *RRTF1* and *RAB18* expression, halotropism and cell damage in *rbohc*, *rhohd* and *rbohf* mutants. Interestingly, *rbohc* and *rbohf* mutant roots exhibited reduced *RAB18* expression compared to wildtype during control and NaCl treatments (Fig. 4C), while *RRTF1* expression was higher in these mutants during salt stress. This aligns well with our observation that ABA responses negatively correlate with sodium-induced responses. The *rbohc* and *rbohf* mutants also showed increased areas of damaged cells during NaCl stress (Fig. 4D/E) and the halotropic response was reduced in these mutants (Fig. S8). Taken together, our data show that sodium-induced responses are repressed by ROS and ABA and are independent of JA accumulation.

**Figure 4.**
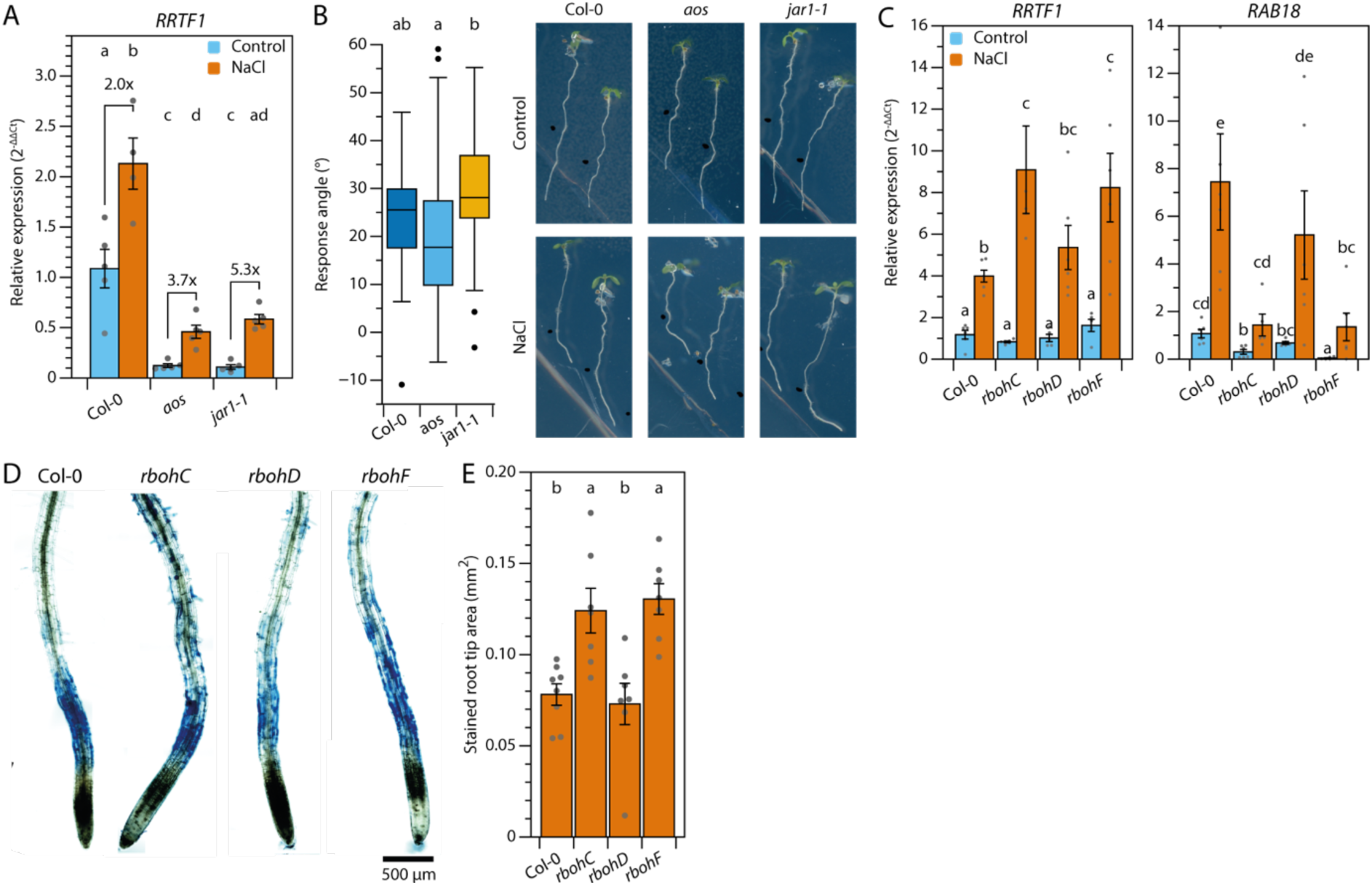
Sodium-induced responses require ROS production but not JA signaling. (A) qPCR analysis of sodium-induced *RRTF1* transcripts in roots of 7-day old JA biosynthesis mutants and Col-0 control. Seedlings were treated with control or 125mM NaCl for one hour. Bars represent mean values +/-SE, n = 5 and individual datapoints are shown as dots. Letters indicate significant differences by two-way ANOVA followed by a Tukey post hoc (p < 0.05). (B) Representative images and quantification of the halotropism response angle of JA signaling mutants (*aos* and *jar1-1*) and wildtype at 48 hours after the introduction of the NaCl gradient (introduced when seedlings were 5-days-old). Different letters indicate significant differences after Dunns test with BH correction for multiple testing (p < 0.05). (C) Relative expression of *RRTF1* and *RAB18* in roots of 7-day old *rboh* mutant seedlings and Col-0 control. Seedlings were treated with control or 125mM NaCl for one hour. Bars represent mean values +/-SE, n = 6 and individual datapoints are shown as dots. Letters indicate significant differences by two-way ANOVA followed by an LSD post hoc (p < 0.05). (D) Representative images of the cell damage assay by Evans Blue staining *rboh* mutants and Col-0 after NaCl-stress for 48 hours. (E) The affected area of the assay (D) was quantified using a script (Fig. S15). Bars represent mean values +/-SE, n=7 - 8 and individual datapoints are shown as dots. Letters indicate significant differences by two-way ANOVA followed by a Tukey post hoc (p < 0.05).

### Sodium-induced gene expression is enhanced in the *snrk2.2/2.3* mutant and is independent of MOCA1 function

To investigate whether sodium-induced gene expression is affected by endogenous ABA signaling, *RRTF1* expression was studied in roots of 7-day-old seedlings in the ABA-insensitive *snrk2.2/2.3* double mutant at 1, 3, 6 and 12 hours after NaCl stress (Fig. 5A/S9) and was found to be significantly higher in NaCl-treated *snrk2.2/2.3* mutants compared to Col-0 from 3 hours onwards. Thus, endogenous ABA signaling can repress the sodium-induced marker gene RRTF1. As expected, the ABA signaling marker gene *RAB18* showed strongly reduced induction by salt in the *snrk2.2/2.3* mutant compared to Col-0 and peaked at 3 hours after stress induction in the wildtype (Fig. S9) ^17,28^.

**Figure 5.**
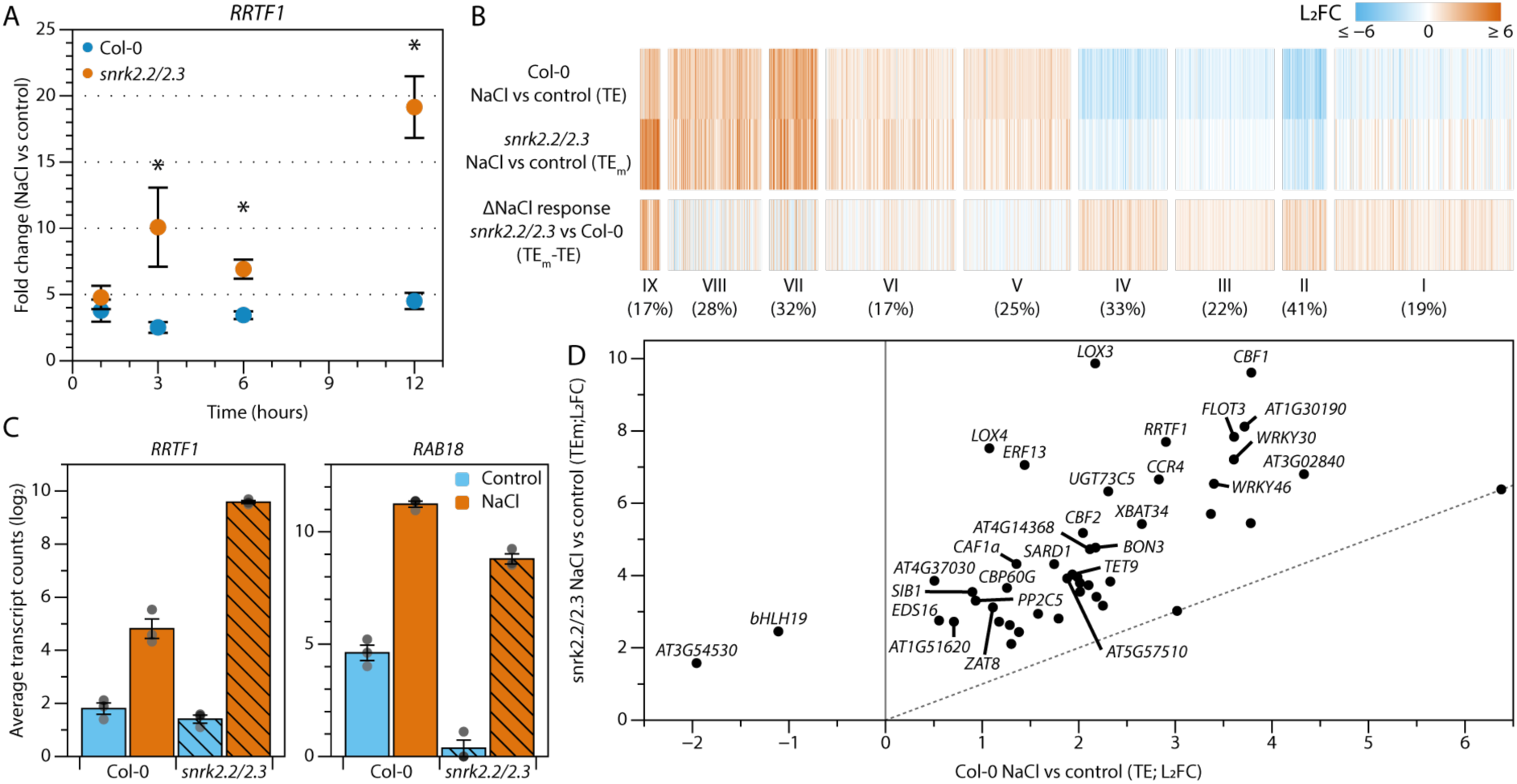
Endogenous ABA signaling is essential for the repression of sodium-induced gene expression. (A) Relative gene expression (Fold change NaCl vs control) of *RRTF1* in the *snrk2.2/2.3* mutant and Col-0. 7-day-old seedlings were transferred to control or 125mM NaCl medium and harvested after 1, 3, 6 and 12 hours. Dots represent the mean (n = 3 - 4) +/- SE and individual datapoints are shown as dots. Data was statistically analyzed using two-way ANOVA per timepoint, followed by a Tukey post-hoc test. Asterisks indicate significant differences between genotypes (p < 0.05). (B) Heatmap of the 1911 overlapping genes that were differentially regulated by NaCl in Col-0 and had a differential response in *snrk2.2/2.3* compared to Col-0 (FDR < 0.05). The numbers below the cluster number represent the percentage of genes overlapping genes between both RNA sequencing experiments. 1161 genes were enhanced or suppressed in *snrk2.2/2.3*xNaCl IT, but not differentially expressed in the Fig. 1 and therefore excluded from this visualization (See Fig. S10C for the analysis including these genes). (C) Normalized counts of *RRTF1* and *RAB18* in Col-0 and *snrk2.2/2.3* at 3 hours. Bars represent averages +/- SE (n=3). (D) Sodium induction of genes in cluster IX (Panel B) in Col-0 versus *snrk2.2/2.3*. The name of genes with a L2FC difference > 2 between Col-0 and *snrk2.2/2.3* are labelled. The dashed line indicates an equal regulation between Col-0 and *snrk2.2/2.3* (L2FC – 0).

**Figure 6.**
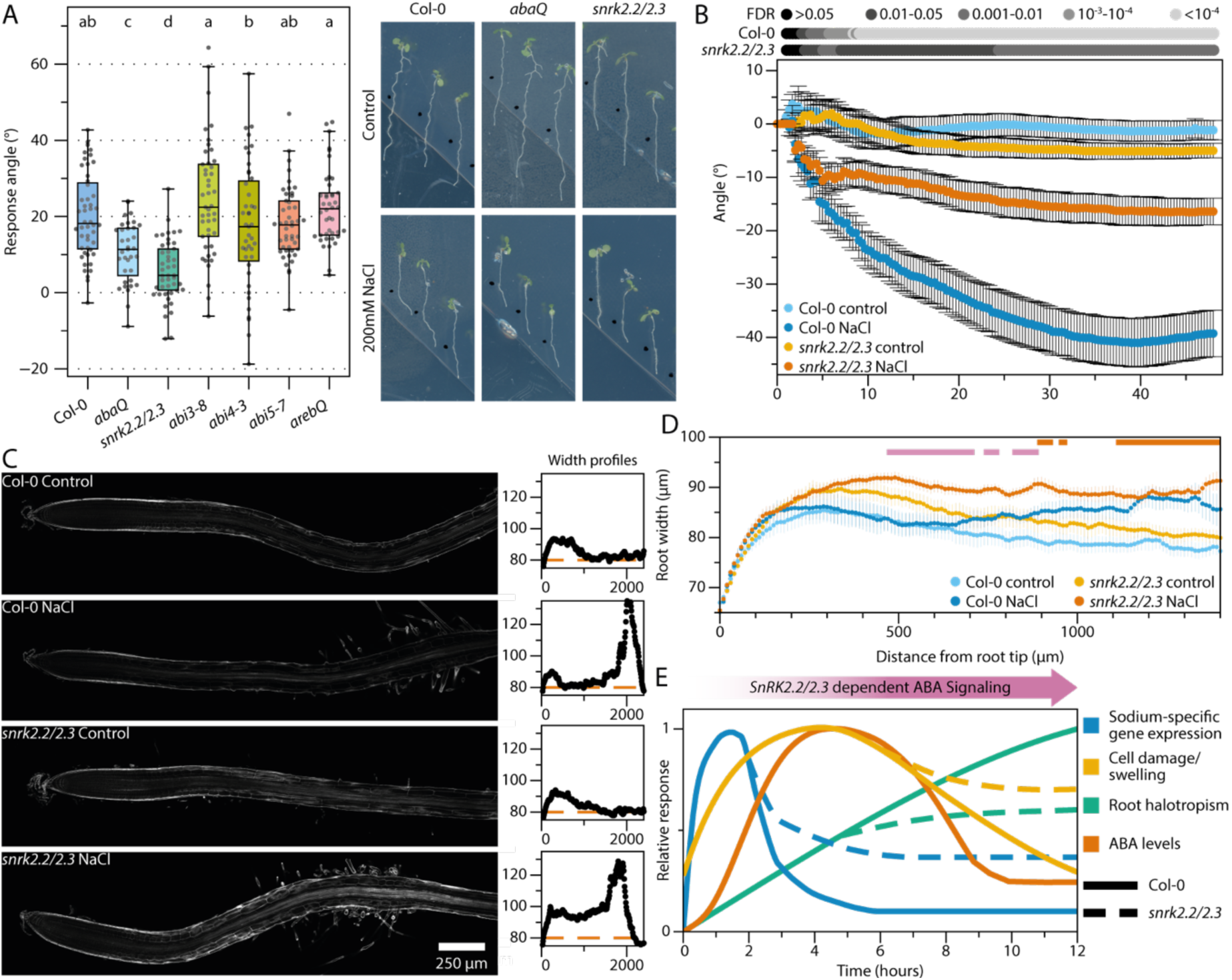
ABA signaling is essential for an adequate salt-specific response of roots. (A) Quantification of the halotropism response of ABA signaling mutants at 48 hours after the introduction of the NaCl gradient. The response angle shows the difference between control and NaCl conditions for the respective mutant. n = 35 - 48 and individual datapoints are shown as dots. Different letters indicate significant differences (p < 0.05), calculated by non-parametrical Dunn’s test and corrected for multiple testing using Benjamin-Hochberg (BH) procedures. Representative images of Col-0, *abaQ* and *snrk2.2/2.3* are shown (see Fig. 13A for the other images). (B) Halotropism time-series with 5-day-old seedlings of *snrk2.2/2.2* double mutants and Col-0 from the moment that the sodium gradient is created. Seedlings were imaged every 20 minutes for 48 hours. Dots represent mean values (n = 20) +/-SE. Pairwise Welch tests were performed for NaCl treated samples and the control condition of the same genotype and corrected for multiple testing with BH. Corrected p-values (NaCl vs control) are shown in grey-values above the graph. (C) Confocal images of calcofluor-white stained Col-0 and *snrk2.2/2.3* roots at 24 hours after NaCl treatment. The width profiles of the depicted roots are shown on the left, with the distance from the root tip (µm) on the x-axis and the width (µm) on the y-axis. The orange dotted line indicates average root width under control conditions. (D) Quantification of root width ratio along the root axis (length 0 = root tip) for Col-0 and *snrk2.2/2.3* as shown as in Panel C. The orange bars above the graph indicate significant differences (FDR < 0.05) between NaCl treated *snrk2.2/2.3* roots and control as analyzed using multiple Welch tests followed by BH correction for multiple testing. The pink bars indicate significance differences between salt-treated Col-0 and *snrk2.2/2.3* roots. Col-0 roots were not significantly different between NaCl and control. (E) A schematic summary of the results presented in this paper. During NaCl stress, sodium ions quickly trigger sodium-induced gene expression, wounding/cell swelling and negative root halotropism. The osmotic component of salt stress triggers the ABA accumulation and signaling and after >3 hours ABA represses sodium-induced gene expression and cell swelling in a *SnRK2.2/2.3*-dependent manner. Therefore, these responses are enhanced in the *snrk2.2/2.3* mutant. Root halotropism is initially ABA-independent but requires ABA signaling to reach full potential.

To obtain a more comprehensive overview of the role of endogenous ABA signaling in sodium-induced gene expression during the response of roots to high salinity, we analyzed the transcriptomic response of the *snrk2.2/2.3* mutant in response to salt after 1 and 3 hours, which are the consecutive timepoints at which the highest NaCl-induced RRTF1 expression was detected (1 hour; Fig. 2A) and where *snrk2.2/2.3* mutant shows a large difference (3 hours; Fig. 5A) with wild-type.

First, we selected genes that had a (1) differential (FDR < 0.05) NaCl response (NaCl vs control) in *snrk2.2/2.3* compared to Col-0 (also termed Interaction Term; Fig. S10A) and (2) were differentially regulated by NaCl in Col-0. We identified 180 and 3072 genes that met both criteria at 1 and 3 hours after stress induction, respectively (Fig. S10B), which aligns well with the relatively slow offset of ABA response (Fig. 2B/C and 5A). Next, we analyzed the 3-hour timepoint in more detail by grouping these selected genes using the clusters as identified in our first transcriptomics experiment (Fig. 1/S10C). The treatment effect (TE; NaCl vs control) in Col-0 showed a highly similar pattern as observed in Fig 1B, in which cluster I-IV and cluster V-IX were down- and upregulated, respectively (Fig. 5B). The genes in clusters I-VIII generally showed reduced salt-induced changes in expression in *snrk2.2/2.3* compared to wildtype (Fig. 5C/D). Strikingly, genes in the sodium-induced cluster IX on the other hand, had a strongly enhanced response in *snrk2.2/2.3*. In other words, these salt-upregulated genes were even higher upregulated by salt in *snrk2.2/3.3* (Fig. 5D). This confirms that endogenous ABA robustly represses sodium-induced genes, a phenomenon that is not limited to the selected marker gene *RRTF1*. To investigate whether the previously reported monovalent cation sensor MOCA1 would be involved in the observed sodium-induced responses, we had included the *moca1* mutant^29^ in our RNA sequencing experiment as well. Only 52 genes showed a differential response to NaCl in *moca1* vs. wildtype (FDR < 0.05; Fig. S11A). No difference was found in NaCl-induced *RRTF1* expression in this mutant compared to wildtype (Fig. S11B).

Interestingly, *moca1* mutants are also not impaired in halotropism, but instead show a hypersensitive root halotropic response, similar to *salt-overly-sensitive* (*sos*) mutants^7^ (Fig. S11C). In conclusion, the sodium-induced responses identified in this paper are not downstream of the monovalent cation sensor MOCA1 and are likely induced by an independent sodium-perception mechanism.

### Endogenous ABA is required to mitigate sodium stress

As endogenous ABA signaling repressed sodium-induced gene expression, we investigated whether it also affects other root responses to sodium stress. First, we studied halotropism in various mutants with defects in ABA signaling: the *pyr1/pyl1/2/4* ABA receptor mutant (referred to as *abaQ*), the *snrk2.2/2.3* protein kinase mutant and mutants for downstream positive regulators of ABA signaling: *abi3-8, abi4-3*, *abi5-7* and *areb1/areb2/abf3/abf1-1* (referred to as *arebQ*; Fig. S1). Halotropism was significantly reduced in both *abaQ* and *snrk2.2/2.3* mutants, showing that core ABA signaling is essential for root directional avoidance of sodium (Fig. 6A/S12A). All investigated mutants for downstream ABA signaling components on the other hand, showed a wildtype halotropic response. This could indicate genetic redundancy in downstream ABA signaling or the involvement of other downstream targets that were not tested here.

To study the temporal characteristics of halotropism in the *snrk2.2/2.3* mutant, 5-day-old plants were placed in a timelapse imaging setup for 48 hours after the introduction of the NaCl gradient (Fig. 6B). Both Col-0 and snrk2.2/2.3 seedlings quickly changed the growth direction of their main root to avoid high sodium concentrations and were significantly different (FDR < 0.05) from control conditions after 2:40 hours and 3:00 hours, respectively. However, the halotropism response stagnated in the *snrk2.2/2.3* mutant after approximately 5 hours. Even though roots of the *snrk2.2/2.3* mutant grew slower than wildtype roots (Fig. S12B), plotting root growth vs angle shows that this could not explain the difference in root angle response between the genotypes. (Fig. S12C). Taken together, these results show that the initial sodium-avoidance response is independent of ABA and ABA is required to enhance the avoidance response at later timepoints. This is consistent with the timing of ABA accumulation and signaling (Fig. 2) and the lack of halotropic response after ABA pretreatment (Fig. 3B).

Previous work reported NaCl-induced cell swelling in the early elongation zone, which is not observed by a comparable osmotic treatment^30^ (and confirmed here in Fig. S13). Interestingly, we observed sodium induced cell damage and *RRTF1* gene expression in this zone (Fig. 3) and halotropism is driven by asymmetrical cell elongation in this zone^7^. Hence, we asked if endogenous ABA signaling also affects cell morphology during salt stress, which we investigated by measuring root width at 24 hours after salt stress induction (Fig. S14A-C). The initial root swelling (1400-2300 µm zone) was comparable between Col-0 and *snrk2.2/2.3* (Fig. S14D). However, salt-treated *snrk2.2/2.3* roots showed significantly more cell swelling compared to control grown seedlings in the region closer to the root tip (900-1500 µm zone) and salt-treated *snrk2.2/2.3* roots were significantly wider than salt-treated Col-0 roots (at 500-900 µm from the tip; Fig. 6C/D), while salt-treated Col-0 roots were not significantly different in width compared to control treated roots. Taken together, this shows that endogenous ABA signaling is important for roots to recover from sodium-induced stress on the transcriptional and cellular level.

## Discussion

In this study, we report that in response to high salt, roots exhibit quick sodium-induced transcriptional responses (1-hour post-stress induction), which are repressed by ABA, which naturally accumulates in response to salinity at later timepoints. Modifying this natural course of action by means of an ABA pretreatment abolishes sodium-induced responses altogether. Moreover, genetic inhibition of ABA signaling during salt treatment leads to (1) increased and prolonged sodium-induced gene expression (Fig. 5), (2) increased root cell swelling (Fig. 6C/D) and (3) reduced negative root halotropism (Fig. 6E). Our data thus reveal an essential role for ABA in mitigating the effects of sodium stress. Sodium-induced gene expression is quickly induced by salt stress but is restored to control levels even in the continuing presence of salt stress at later timepoints (Fig. 3). We found that these responses are enhanced and prolonged in the ABA-insensitive mutants and on the other hand abolished altogether when applying an ABA pretreatment, which confirms that the restoration to normal root functioning after root response to high salinity depends on ABA signaling. The halotropism response is more complex as it is reduced by either ABA pretreatment or genetic ABA insensitivity (Fig. 3/6). This may seem contradictive, but as redirection of root growth depends on an asymmetry in cell elongation^31,32^, we hypothesize that disturbing this asymmetry either by genetically blocking or treating with ABA homogeneously, hampers the halotropic response. Interestingly, the *snrk2.2/2.3* halotropism timeseries data suggest that *snrk2.2/2.3* mutants can initially redirect root growth to avoid high salt concentrations like wildtype, but the response stagnates after 5:00 hours (Fig. 6B). This aligns well with the halotropism phenotype after the ABA pretreatment, in which roots initially bend, but no net root avoidance is measured after 24 hours. Thus, while not essential for the initial redirection root growth, ABA could be required to inhibit gravitropism and thus maintenance of the sodium avoiding halotropic response at later timepoints. Notably, *rbohc* and *rbohf*, but not *rbohd* mutants, also show reduced ABA marker gene (*RAB18*) expression under control and salt treatment, and enhanced sodium-induced gene expression and reduced halotropism. These mutants thus phenocopy *snrk2.2/2.3* with respect to sodium-induced responses of the root. This aligns well with existing literature which shows that responses to ABA (root growth and stomatal closure) are reduced in *rbohf*, but not in *rbohd*^33^ mutants. Therefore, RBOHF-mediated ROS production is recognized as a secondary messenger to ABA signaling^33^.

The quick induction and consequent repression of sodium-induced gene expression is highly correlated with previously reported root growth phases during early salt stress^17^. After exposure to NaCl, roots go through a stop (0-5 hours), quiescent (5-9 hours), recovery (9 – 12 hours) and homeostasis (>12 hours) growth phase^17,34^. Our timeseries show that *RRTF1* expression is high at the early timepoints after stress induction (Fig. 2, Fig. 5A), which corresponds with the stop and quiescent phase. Later, the osmotic stress-induced ABA biosynthesis and signaling repress sodium-induced gene expression, which aligns with the timing of growth recovery. These correlations suggest that increased expression of sodium-induced genes causes growth reduction. Supporting this hypothesis, overexpression of some highly expressed sodium-induced genes (*RRTF1*^24,35^, *DWARF AND DELAYED FLOWERING 1* (*DDF1*)^36^ or *CALMODULIN BINDING PROTEIN 60G* (*CBP60g*)^37^) leads to severe dwarfism. In this model, ABA signaling (and thus, the repression of sodium-induced genes) is required to restore root growth. Indeed, chemical inhibition of ABA biosynthesis during salt stress inhibits growth recovery after the quiescent phase^17^. Taken together, this indicates that sodium-induced gene expression correlates with early root growth dynamics after salt stress exposure.

To get a better understanding of sodium-induced regulatory mechanisms, we aimed to disentangle the sodium-induced, general monovalent cation, and osmotic responses to salt stress. We expected to identify monovalent-cation specific transcriptional responses, as salt stress specifically causes a rapid (<20 seconds) increase in intracellular Ca^2+^ which migrates in waves throughout the plant. These waves cause transcriptional changes in the shoot, which is not directly in contact with the saline soil^19^. Recently, it was described that these Ca^2+^ waves are not sodium-induced, but are induced by other monovalent cations (Li^+^ and K^+^) as well, and are sensed by negatively charged sphingolipids (biosynthesized by MOCA1^29^), which in turn activate yet unknown Ca^2+^ channels^29^. In our transcriptome analysis we were able to distinguish sodium-induced gene expression from a general osmotic stress response induced by both KCl and osmotic stress, as well as ABA treatment. However, no evident generic monovalent cation stress pattern was identified, which aligns well with the almost wildtype-like salt stress transcriptome of the monovalent cation sensor mutant *moca1*. In accordance, we found MOCA1 not to be required for the sodium-induced halotropism response (Fig. S11). This suggests that monovalent cation-induced Ca^2+^ waves do not affect early sodium-induced responses in the roots in our conditions, leaving open the question of the molecular nature of sodium perception in plants.

We used *RRTF1*, a strong sodium-induced transcript as a marker gene to characterize the regulatory mechanisms of sodium-induced gene expression. Previously, we showed that *RRTF1* expression is downstream of MAP KINASE6 (MPK6)^34^, but up- and downstream phosphorylation cascades remain unknown. We investigated a possible role for JA because elevated sodium levels cause cell damage and *RRTF1* expression is strongly induced by damage in a JA-dependent manner^22–24^. However, while JA mutants have reduced *RRTF1* expression under control conditions, the expression was still induced by salt. Genetic explorations of other signaling pathways, including ROS production (*rbohc/f*, Fig. 4C-E), ABA (*snrk2.2/2.3,* Fig. 5/6) and cell wall maintenance (*fer-4* and *herk1 the1-4* ^34^) all showed enhanced expression of *RRTF1*. Because *RRTF1* expression is induced by multiple factors, we hypothesize that these factors regulate cellular functioning during salt stress, and misregulation therefore makes plants more susceptible to salt stress.

To investigate the cause of these enhanced stress responses, we focused on the biological processes associated with sodium-induced responses. Interestingly, the sodium-induced gene cluster (Fig. 1) and genes with enhanced expression in *snrk2.2/2.3* (Fig. 5) are associated with biotic stress/wounding responses. It was shown that these processes are also differentially regulated in the transcriptome of salt-stressed *fer-4*, which has increased salt-induced cell swelling (and even bursting)^30^. Strikingly, despite being more subtle than *fer-4* mutants, *snrk2.2/2.3* mutants also showed increased cell swelling in the apical root (Fig. 6). This aligns well with previous reports in which NaCl-induced cell swelling is strongly enhanced when ABA biosynthesis is chemically inhibited^17^. Nevertheless, even though *fer-4* and *snrk2.2/2.3* both have a cell swelling phenotype, ABA signaling is enhanced in *fer-4* mutants^38,39^, highlighting again the complexity of regulation of cell wall modifications during salt stress in roots^30,34^. On the other hand, ABA biosynthesis and signaling mutants also have a strongly reduced cellulose content^40,41^ and inhibition of cellulose biosynthesis causes hypersensitivity during salt stress^30,42^. Furthermore, ABA-mediated alterations in microtubule organization during salt stress are important to strengthen the cell wall and limit cell damage^31^. Taken together, we hypothesize that weakened cell walls in *snrk2.2/2.3* roots could contribute to the observed increased cell swelling induced by NaCl.

To conclude, we have shown that when plant roots are exposed to salinity, early sodium-induced gene expression is repressed by the slower osmotic stress-induced ABA accumulation and signaling. By enhancing (ABA treatment) or repressing (ABA-insensitive mutants) ABA signaling, we were able to show that the interaction between the stress components is required to mount an adequate response to salt while maintaining cell integrity. Our work gives novel insights in the tight temporal and spatial responses of a complex stress. We argue that salinity stress, like many abiotic stresses, triggers different response pathways, which are separated in time and space. The existence of a clearly specific response opens new doors to investigate sodium sensing mechanisms and targets for salt stress resilience.

## Acknowledgements

We thank Zhen-Ming Pei for providing *moca1* seeds, Nora Gigli-Bisceglia for providing *aos* and *jar1-1* seeds, Elwira Smakowska-Luzan for providing *rbohd* seeds, and Scott Hayes for providing *abaQ* and *arebQ* seeds.

## Author contributions

Conceptualization: J.L., Y.Z., C.M.M.G., C.T.; Experimentation: J.L, Y.Z, E.v.Z, A.J.M, T.d.Z, F.V, M.V., A.O.D-A; Software: J.L.; Formal analysis: J.L.; Writing - original draft: J.L., C.M.M.G. ,C.T.; Writing - review & editing: J.L, Y.Z, E.v.Z, A.J.M, T.d.Z, F.V, M.V., A.O.D-A, C.M.M.G., C.T.;

Visualization: J.L..; Supervision: C.T, C.M.M.G.; Project administration: C.T.,Y.Z.; Funding acquisition: C.T., Y.Z.

## Data availability

Raw and processed RNA sequencing data files are available via ArrayExpress (E-MTAB-13345 for Fig.1 RNAseq), or to be made available via ArrayExpress upon publication (Fig. 5 RNAseq).

## Online materials and methods

### Plant materials, growth conditions and stress assays

*Arabidopsis thaliana* plants of ecotype Columbia-0 (Col-0) were used in all experiments. Previously described lines were used here *aos*^43^, *jar1-1*^44^, *rbohc*(SALK_071801)^45^, *rbohd-1* (SALK_070610C)^46^, *rbohd-2* (SALK_120299C)^47^, *rbohd-3*^48^*, rbohf-1* (SALK_059888)^49^, *rbohf-2* (SALK_034674)^27^, *snrk2.2/2.3* (GABI_807G04/ SALK_107315)^14^, *abi3-8*^50^, *abi4-3*^50^, *abi5-7*^50^, *abaQ* (*pyr1-1*/*pyl1-1*/*pyl2-1*/*pyl4-1;* Point/Salk_054640/CSHL_GT2864_1/Sail_517 _C08)^12^, arebQ^51^, (*areb1*/*areb2*/*abf3*/*abf1-1;* SALK_002984/SALK_069523/SALK_096965/ SALK_132819), *pRRTF1:RRTF1-VENUS* (Named; *ERF109pro:ERF109-Venus*)^22^, *moca1* ^29^. Seeds were wet sterilized by 30% commercial bleach and 0.2% (v/v) triton-X 100 for 10 minutes and washed 5 times with sterile milliQ. Seeds were sown on ½ Murashige and Skoog (MS) medium including vitamins (Duchefa) containing 0.1% 2-Morpholinoethanesulfonic acid monohydrate (MES) buffer (Duchefa) and 1% Daishin agar (Duchefa) and pH was adjusted to 5.8 with KOH. Seeds were stratified at 4 °C in the dark for 2 days. Plants were grown in a rack at an angle of 90° at 22°C and long day photoperiod (16 hours light, 120µM/m^2^/s).

### Pharmacological treatments and root harvesting

Agar plates were covered with 1.5×10.5cm 50µM nylon mesh strips (Sefar BV) before sowing to facilitate seedling transfer to treatment plates. Plants were grown for 7 days (6.5 for the 12-hour ABA (5mM stock in ethanol) pretreatment) under the conditions described above and transferred to treatment plates. Roots were dissected with a sharp blade and flash frozen in liquid nitrogen. For RNA extraction and ion measurement, approximately 60 roots were pooled in one sample.

For the first RNA sequencing experiment (Fig. 1), stress induction was performed as described above with minor changes. Instead of small mesh strips, the whole agar plate was covered by a 10.5×10.5cm 50µM nylon mesh. Plants were grown for 8 days at a 70° angle before transfer to 130 mM NaCl (Duchefa), 130 mM NaNO_3_ (Merck), 130mM KCl (Merck), 260 mM Sorbitol (Duchefa), 30mM LiCl (Merck), 25µM ABA (Sigma, in ethanol) or control medium without supplements. Roots were harvested after 6 and 24 hours. Approximately 40 roots from one plate were pooled as one biological replicate.

For the second RNA sequencing experiment (Fig. 5), plants were grown and harvested in the same way, with the only exception that 0.8% Plant agar (Duchefa) was used instead of Daishin agar.

### RNA isolation and qPCR

Samples were homogenized using 2 stainless steel beads and Mixer Mill MM400 (Retsch), after which RNA was isolated with a Total RNA Isolation kit (NZYTech)^52^. RNA quality was quantified using the Nanodrop™ One, followed by RQ1 DNase treatment (Promega) with 2000ng RNA. 1000ng was used for cDNA synthesis using iScript reverse transcriptase (Bio-Rad). cDNA was diluted to 10ng/µL before quantification by RT-qPCR (25ng cDNA per sample), using SYBR Green blue mix lo-rox (Sopachem) with a CFX96 real time system (Bio-Rad). Relative expression was calculated with the ΔΔCt method, using *MON1* (AT2G28390) as housekeeping gene. Primers used for RT-qPCR are listed in Table S7.

Samples for RNA sequencing were isolated using the RNeasy Kit (Qiagen). For the first RNA sequencing in combination with Tripure (Roche) as previously published^52^.

### Library preparation, sequencing and data processing

Total RNA was used for RNA library preparation suitable for Illumina HiSeq paired end sequencing using Illumina’s TruSeq stranded RNA sample prep kit using polyA mRNA selection. mRNA was further processed directly including RNA fragmentation, first and second strand cDNA synthesis, adapter ligation and final library amplification following manufacturers protocol. The final library was eluted in 30 µl elution buffer followed by quality assessment using a Bioanalyzer 2100 DNA1000 chip (Agilent Technologies) and quantified on a Qubit quantitation platform (Life Technologies).

Prepared libraries were pooled and diluted to 6 pM for TruSeq Paired End v4 DNA clustering on one single flow cell lane using a cBot device (Illumina). Final sequencing was done on an Illumina HiSeq 2500 platform using 126, 7, 126 flow cycles for sequencing paired end reads plus indexes reads. All steps for clustering and subsequent sequencing were carried out according to the manufacturer’s protocol. Reads were split per sample by using CASAVA 1.8 software (Illumina Inc, San Diego CA, USA). All sample preparations and sequencing were done by the Genomics lab of Wageningen University and Research, Business Unit Bioscience.

RNA poly-A enrichment library preparation and transcriptome sequencing (150bp paired-end mode) on a NovaSeq 6000 platform of the second RNA sequencing experiment were conducted by Novogene UK Co. Ltd (Cambridge, UK).

First read quality was analyzed with FastQC^53^ and MultiQC^54^ packages in Python 2.7, followed by trimming of low quality reads with Trim Galore!^55^. The reads were re-examined with FastQC and MultiQC after trimming and mapped to the Arabidopsis TAIR10 transcriptome^56^ using Salmon^57^. DESeq2^58^ package in R^59^ was used for differential gene expression analysis. Both timepoints were imported and analyzed separately in DESeq2. Heatmaps were created with the R package pheatmap^60^, using Euclidean distance mapping and Ward.D clustering algorithms. GOseq^61^ was used for Gene Ontology (GO) enrichment analysis using GO_SLIM annotations from TAIR (version 2021-10-01). Enrichment was calculated using Fishers exact test with Benjamin-Hochberg correction for multiple testing.

### Ionomics

Roots were harvested at 6 hours after treatment and washed 3 times in milliQ water. The harvested material was dried for 24 hours at 60 degrees. The samples were measured using inductively coupled plasma mass spectrometer (IPC-MS) by the Nottingham Ionomics Facility (UK). All handling was done using plastic tweezers.

### Halotropism

Growth conditions as described above, but medium contained 0.5% sucrose. After five days, roots were aligned to be 5mm from the cutting edge of the gradient, which was introduced by removing the lower part of the medium and replacing it with either 1/2MS control or 200mM NaCl medium. Plants were scanned at 24 and 48 hours after the introduction of the gradient. ABA (5mM stock in ethanol; Sigma) pretreated roots on a control (no NaCl) treatment had the tendency to grow away from the agar plate (n=7, 15% at 24 hours and n=15, 33% at 48 hours) and were excluded from quantification. This response was not observed for the ABA pretreated seedlings that received the NaCl treatment. After 48 hours, plates were scanned with an Epson Perfection V800 at 400 dpi. Root growth and angle were quantified after 24 hours using Fiji with the SmartRoot plugin^62^. The timeseries was created by an automated infrared imaging setup, which has been described before^34,63^. Plants were grown in this growth chamber and were imaged after the introduction of the gradient. Roots were traced using an automated script (https://github.com/jasperlamers/timelapse-backtracing) as described before^34^.

### Evans Blue staining

Whole seedlings were emerged in an Evans Blue solution (0.25% w/v, Sigma) for 15 minutes. Plants were destained for 15 minutes in sterile milliQ and the apical root was imaged with a DM2500 optical microscope (Leica) using a 10x objective. Images were stitched using pairwise stitching^64^ in the Fiji software^65^. The largest continuous stained area was segmented with an automated script in Python3.7 using OpenCV (v4.5.1.48; Fig. S15). Evans Blue was stained at 24 hours after stress induction for the confocal experiments (excitation 560, emission, 625-760).

### Root width analysis and RRTF1-mVenus imaging

Roots were treated for 2 hours (*pRRTF1::RRTF1-Venus*) or 24 hours (root width) at 125mM NaCl, followed by fixation using 4% PFA and clearing using ClearSee^66^. Roots were stained with CalcoFluorWhite and imaged on Stellaris 5 Confocal LSM (Leica) using a 10x dry lens. (excitation 405nm, emission, 425-475nm). Venus was excited at 515m, and emission was detected bwteen 525-550nm.

### Hormone measurements

Hormone accumulation measurements were performed as described before, following the exact same procedure using root tissue rather than whole seedlings^67^. In short: after 7 days the plants were transferred to fresh agar plates supplemented with 125 mM NaCl (Duchefa), 125 mM KCl (Merck), 250 mM Sorbitol (Duchefa) or ½ MS control medium without supplements. After 1, 3, 6, 12 hours roots were dissected with a sharp blade, weighed, flash frozen in liquid nitrogen and stored at -80°C. 60 roots (11.8±3.0 mg material) were pooled in one biological replicate. Frozen material was ground to a fine powder using 2 stainless steel beads at 50 Hz for 1 minute in a paint shaker (Fast & Fluid). Ground samples were extracted with 1 mL of 10% methanol containing 100nM stable isotope-labeled internal standards for each investigated compound (Supplemental Table 2). Samples were extracted using an existing protocol with modifications^68^. Namely, a StrataX 30mg/3mL spe-column (Phenomenex) was used. Solvents were removed by speed vacuum system (ThermoSavant).

For detection and quantification by liquid chromatography-tandem mass spectroscopy, sample residues were dissolved in 100µL acetonitrile /water (20:80 v/v) and filtered using a 0.2 µm nylon centrifuge spin filter (BGB Analytik). ABA was quantified by comparing retention times and mass transitions with ABA standards using a Waters XevoTQS mass spectrometer equipped with an electrospray ionization source coupled to an Acquity UPLC system (Waters) as previously described^69,70^. Chromatographic separations were conducted using acetonitrile/water (+0.1% formic acid) on a Acquity UPLC BEH C18 column (2.1 mm ×100mm, 1.7µm, Waters) at 40°C with a flowrate of 0.25 mL/min. First the column was equilibrated for 30 minutes using the solvent (acetonitrile /water (20:80 v/v) + 0.1% formic acid). Samples were analyzed by injecting 5µL, followed by the elution using program of 17 minutes in which the acetonitrile fraction linearly increased from 20% (v/v) to 70% (v/v). The column was washed after every sample by increasing the acetonitrile fraction to 100% in one minute and maintaining this concentration for 1 minute. The acetonitrile fraction was reduced to 20% in one minute and maintained at this concentration for one minute before injecting the next sample. The capillary voltage was set at 2.5kV, the source temperature at 150°C and the desolvation temperature at 500°C. Multiple reaction monitoring was used for quantification^69^. Parent-Daughter transitions and cone voltages were set using the IntelliStart MS Console. Peak quantification was processed with Targetlynx. Samples were normalized for the internal standard recovery and sample weight. All values were normalized to the 1-hour control timepoint.

### Statistical analysis

Single timepoint data was analyzed using Levene’s test to verify equal variances (p > 0.05) and Shapiro-Wilk to verify normal distribution (p > 0.05). Two-way ANOVA followed by a post-hoc Tukey test when samples were normally distributed and had an equal variance. Alternatively, samples were analyzed using the non-parametric Dunn test preceded by a Kruskal-Wallis test (p < 0.05). Benjamini-Hochberg was used to correct for multiple testing. Halotropism timeseries and root witdth were analyzed using non-paired t-tests per timepoint or length unit (NaCl vs control per genotype), respectively, and corrected for multiple testing using the Benjamini-Hochberg method. Other multi-timepoint data was analyzed per timepoint, using Two-way ANOVA followed by Tukey post-hoc test (after Levene’s test to verify equal variances (p > 0.01) and Shapiro-Wilk to verify normal distribution (p > 0.05)). All analyses were conducted in R and Microsoft Excel.

### Accession numbers

*RRTF1*, AT4G34410; *RAB18*, AT5G66400; *LTI78*/*RD29A*, AT5G52310; *SnRK2.2*, AT3G50500; *SnRK2.3*, AT5G66880; *PYR1*, AT4G17870; *PYL1*, AT5G46790; *PYL2*, AT2G26040; *PYL4*, AT2G38310, *ABI3*, AT3G24650; *ABI4*, AT2G40220; *ABI5*, AT2G36270; *AREB1*, AT1G45249; *AREB2*, AT3G19290; *AREB3*, AT3G56850; *ABF1*, AT1G49720; *RBOHC*, AT5G51060; *RBOHD*, AT5G47910; *RBOHF*, AT1G64060; *AOS*, AT5G42650; *JAR1*, AT2G46370; *MON1*, AT2G28390, *MOCA1,* AT5G18480.

## Supplemental Figures

**Figure S1.**
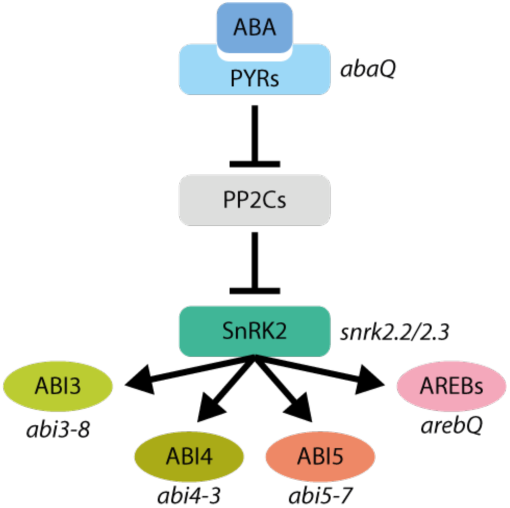
Simplified representation of the ABA signaling pathway, indicating the ABA mutants used in this study. ABA is sensed by *PYRABACTIN RESISTANCE1 (PYR)/PYR1-LIKE* family of receptors, which releases the inhibition by PROTEIN PHOSPHATASES TYPE 2C (PP2C) proteins of the SUCROSE NON-FERMENTING 1-RELATED PROTEIN KINASE2 family (SnRK2.2). SnRK2s activate downstream transcription factors (TFs) such as ABA INSENSITIVE 3 (ABI3), ABI4, ABI5 and ABA-RESPONSIVE ELEMENT-BINDING FACTOR/ABSCISIC ACID RESPONSIVE ELEMENT-BINDING FACTOR 1 TFs (AREB/ABF)

**Figure S2.**
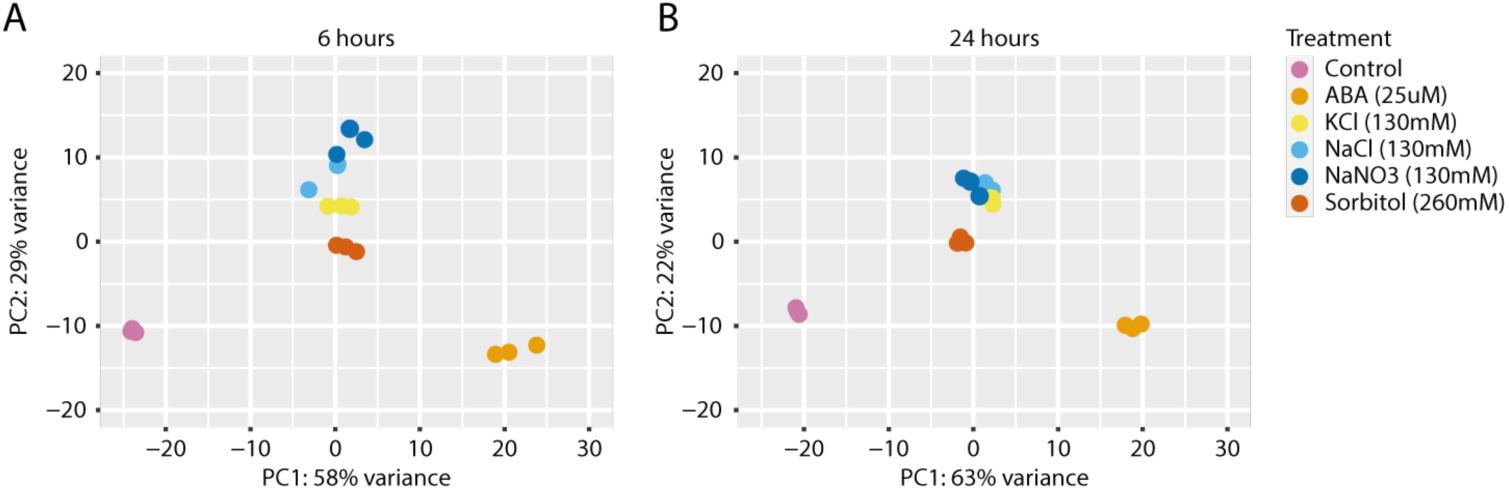
Stress-specific signaling in response to sodium ions is more apparent at 6 hours after treatment. (A-B) Principal component analysis (PCA) for normalized counts of all transcripts of root tissue of 8-day old seedlings treated for 6 or 24 hours. The included treatments were 130mM NaCl, 130mM NaNO_3_, 130mM KCl, 260mM sorbitol and 25µM ABA on solid ½MS medium with 1g/L MES buffer. The first two components are presented. Colors indicate the different treatments. (A) 6 hours and (B) 24 hours after stress initiation.

**Figure S3.**
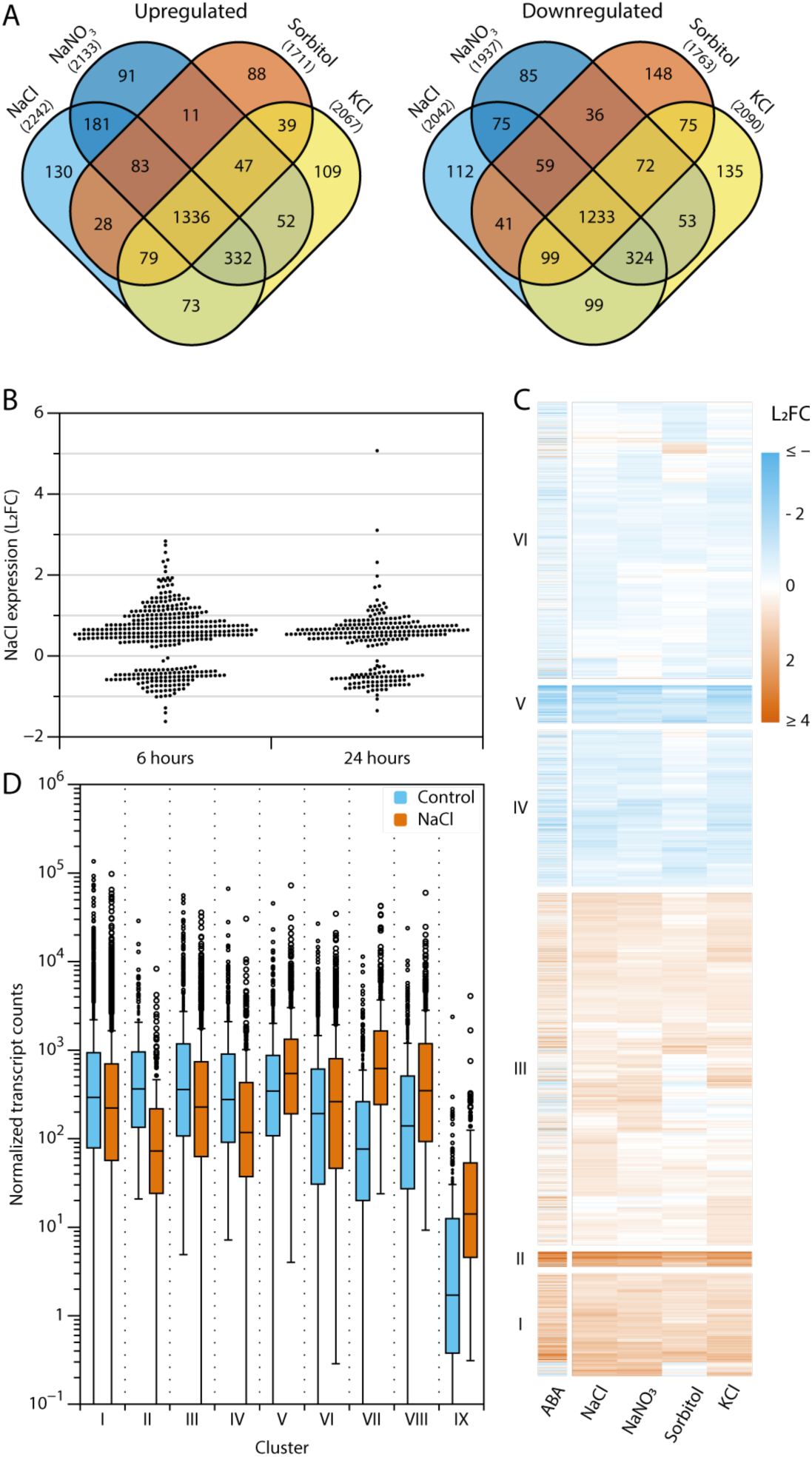
Osmotic and ionic stress and ABA-induced gene expression responses mostly overlap at 24 hours. DEGs were defined based on significance (adjusted p < 0.05) and expression (LFC > |0.5|) and selected when these conditions were met in at least one of the 4 treatments (130mM NaCl, 130mM NaNO_3_, 130mM KCl or 260mM sorbitol). (A) Venn diagrams showing the overlap of differential gene expression between the different treatments vs the ½MS control treatment. (B) The expression of sodium-specific genes at 6 and 24 hours after the NaCl treatment. Sodium-specific genes were identified using the Venn diagrams (Fig 1A and S3A). (C) Heatmap of all 5213 DEGs. Treatments (columns) and genes (rows) were clustered with Euclidean distance mapping and Ward1 clustering. ABA (25µM) was not used for clustering and was added afterwards as an annotation column. Colors indicate the Log_2_FoldChange. The heatmap was divided into 6 clusters. The numbering is indicated below the heatmap. (D) Normalized transcript counts per cluster in control and NaCl-treated samples.

**Figure S4.**
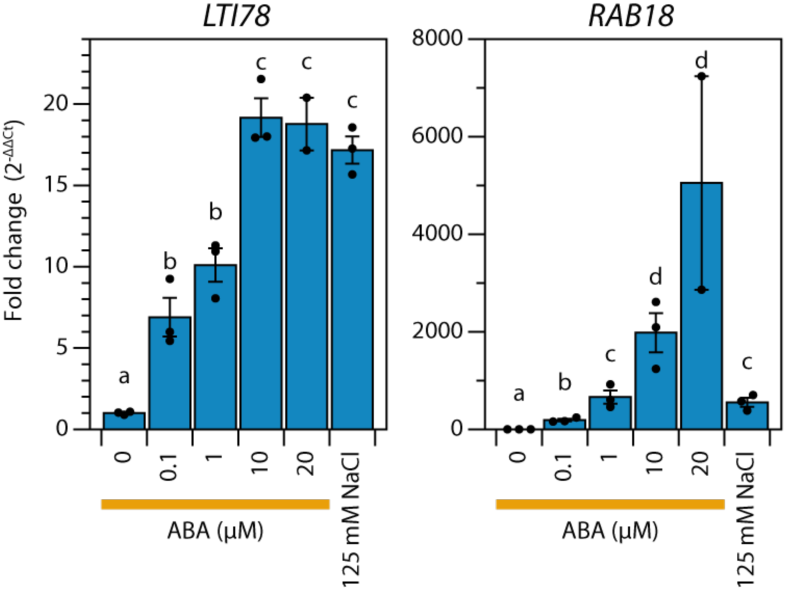
125 mM NaCl-induced ABA marker gene expression corresponds to ABA-induced expression of a 1-10µM ABA treatment. Comparison of ABA-induced and NaCl-induced ABA marker gene expression (*LTI78* and *RAB18*) in 7-day-old Col-0 roots at 6 hours after the application of a range of ABA concentrations, or 125mM NaCl. Letters indicate significance (ANOVA + Tukey HSD, p <0.05). Bars represent the average (n=2-4) +/- SE.

**Figure S5.**
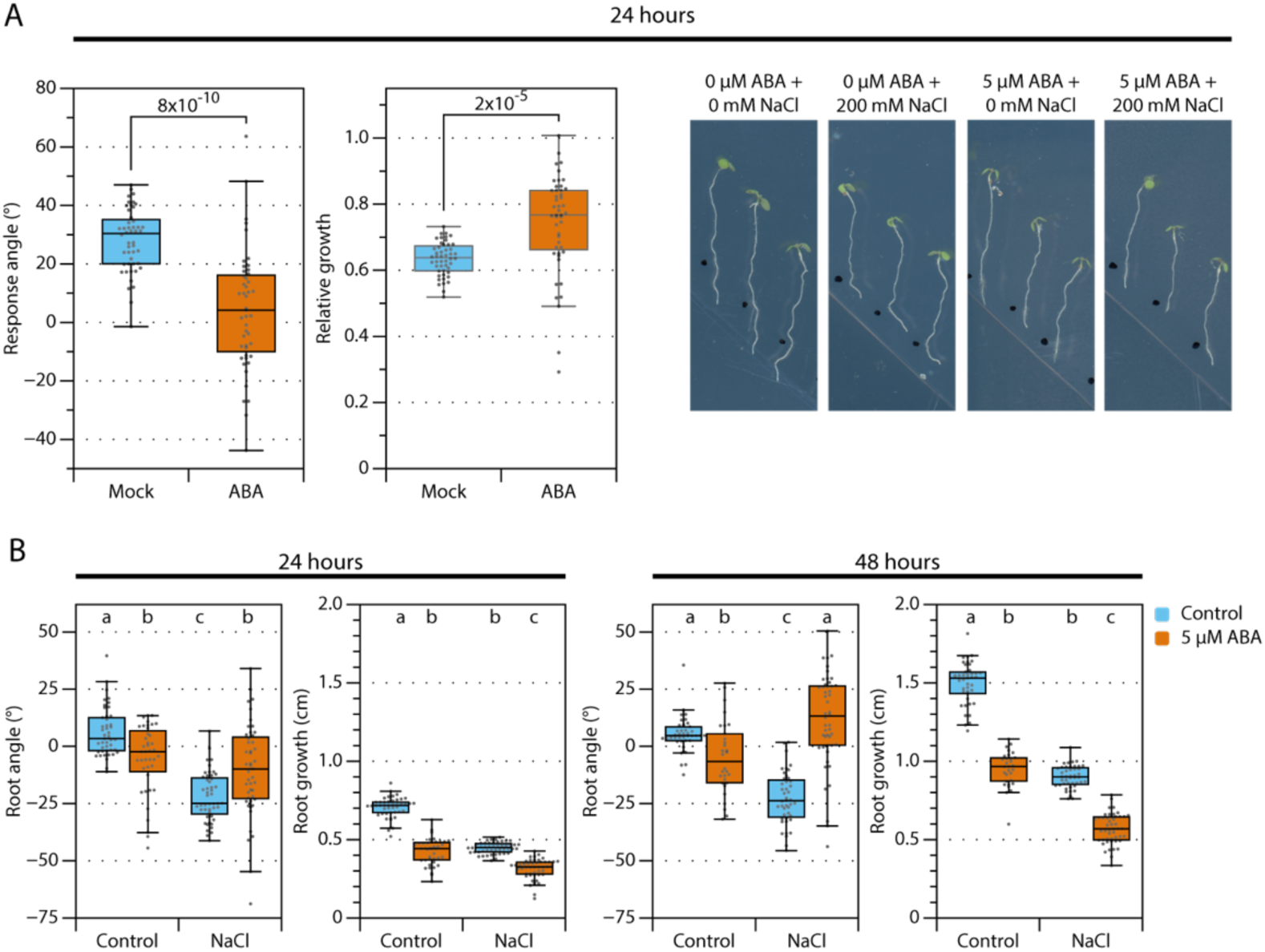
ABA-pretreatment represses halotropism. (A) Representative images and quantification of the halotropism response with or without a 12-hour ABA pretreatment in 5-day-old seedlings at 24 hours after the introduction of the gradient. The relative growth shows the ratio between NaCl and control conditions for the respective ABA pretreatment. Statistics were performed using pairwise t-tests. (B) Absolute data of the halotropism with ABA-pretreatment at 24 and 48 hours after introduction of the gradient (As shown in Fig. 3B). Different letters indicate significant differences after Dunns test with BH correction for multiple testing (p < 0.05). At 24 hours, (n=40-48) and at 48 hours (n=30-48).

**Figure S6.**
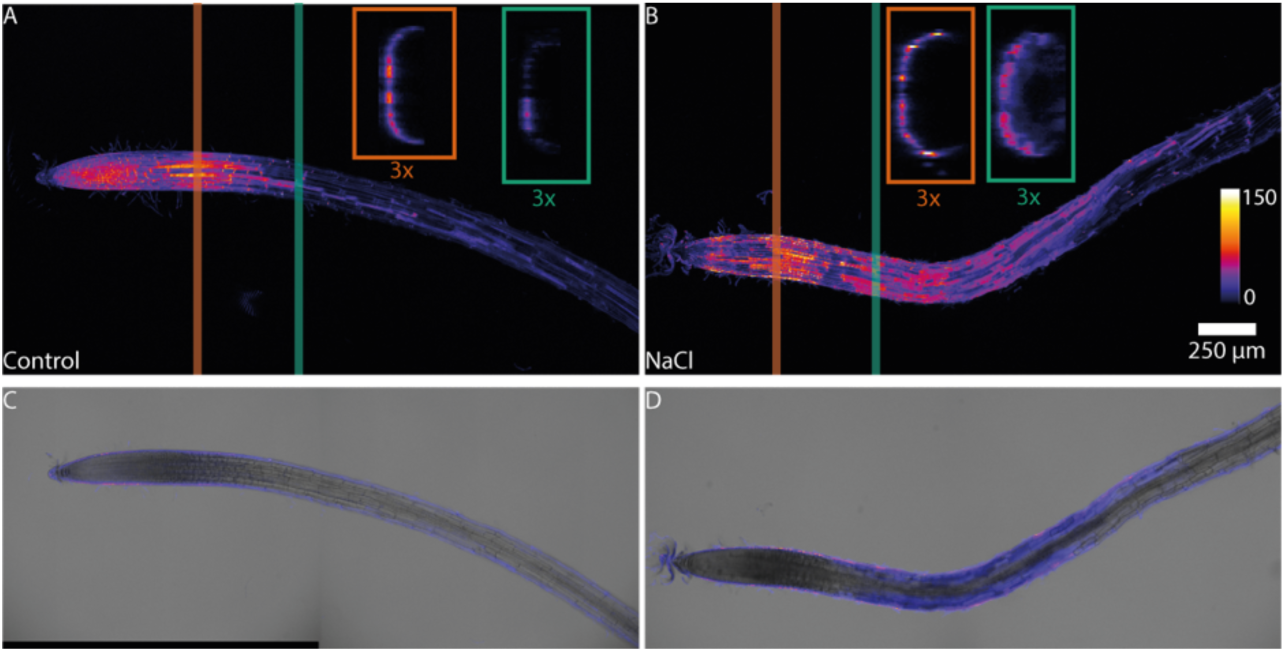
Evans Blue causes a superficial signal in control roots that is distinctive from the NaCl induced cell damage. (A-B) Max projections of the Evans Blue staining in roots treated with (A) control or (B) 125mM NaCl for 24 hours. The lines indicate the location of the 3x magnified orthogonal views (boxes) in the corresponding color. Locations were chosen based on the Evans Blue staining. (C-D) Composites images of Evans Blue staining and the brightfield of the midplane (1 stack) of the roots shown in panel A/B.

**Figure S7.**
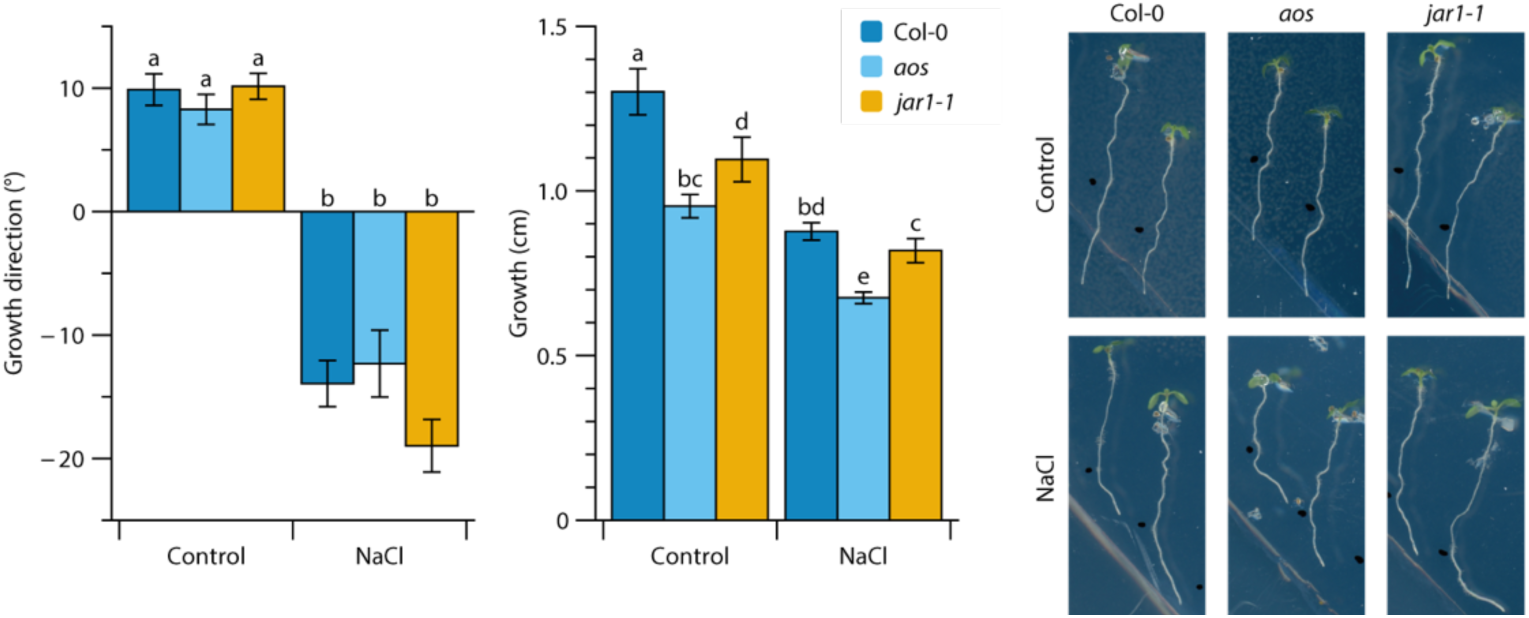
Sodium-specific responses are not JA-dependent. Absolute data of the halotropism assay shown in Fig. 4B. Quantification of the halotropism response angle and relative growth of the main root at 48 hours after the introduction of the NaCl gradient (introduced when seedlings were 5-days-old). JA signaling mutants (*aos* and *jar1-1*) and Col-0 control (n=35). Bars represent averages +/- SE. Different letters indicate significant differences after Dunns test with BH correction for multiple testing (p < 0.05).

**Figure S8.**
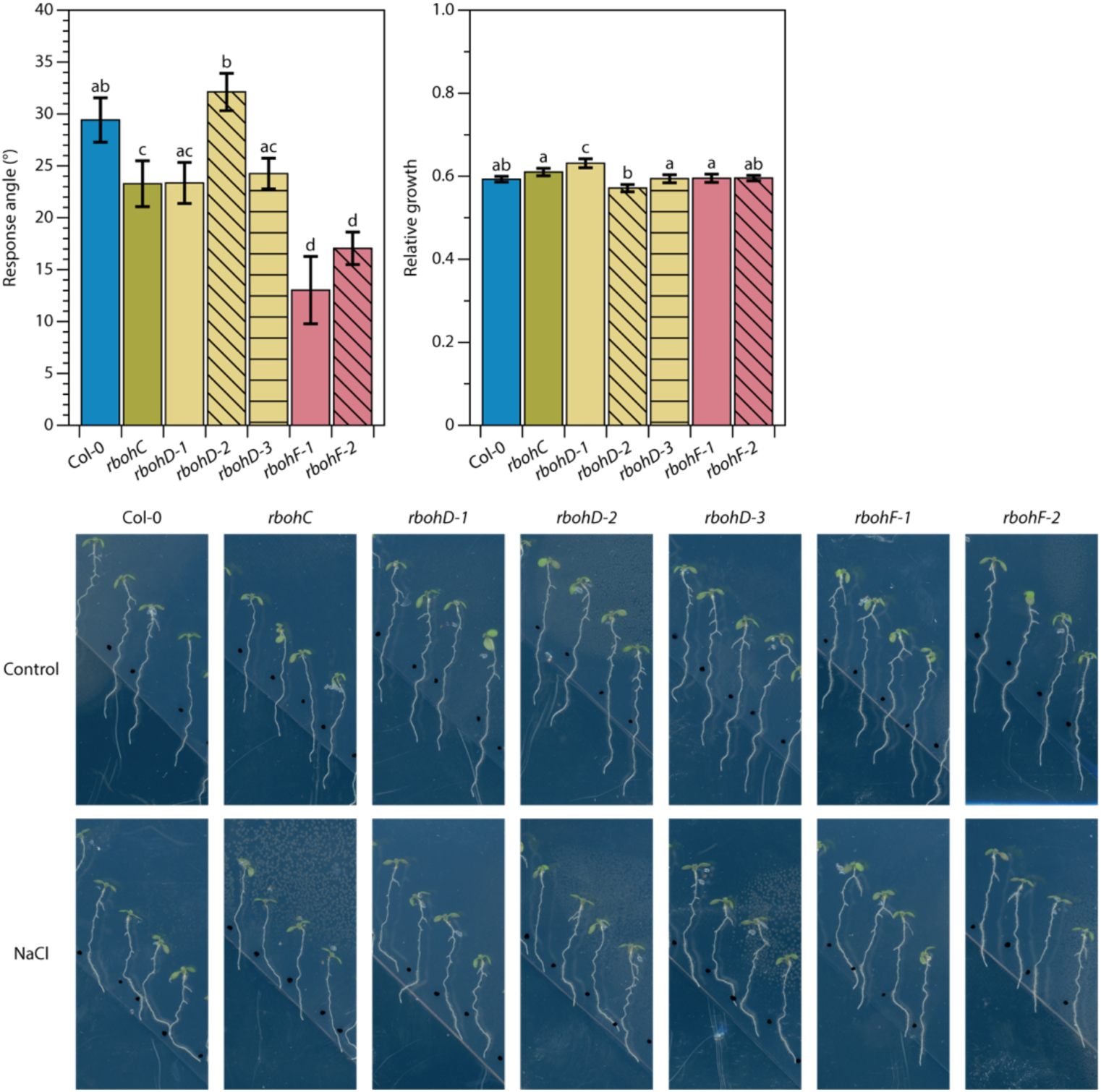
Extracellular ROS production is important for halotropism. Quantification and representative images of the halotropism response angle and relative growth of *rbobc* (1 allele)*, rbobd* (3 alleles) and *rbobf* (2 alleles) mutants and Col-0 control at 48 hours after the introduction of the NaCl gradient (introduced when seedlings were 5-day-old, n=42-45). Bars represent averages +/- SE. Letters indicate significant differences after Dunn’s test (p < 0.05).

**Figure S9.**
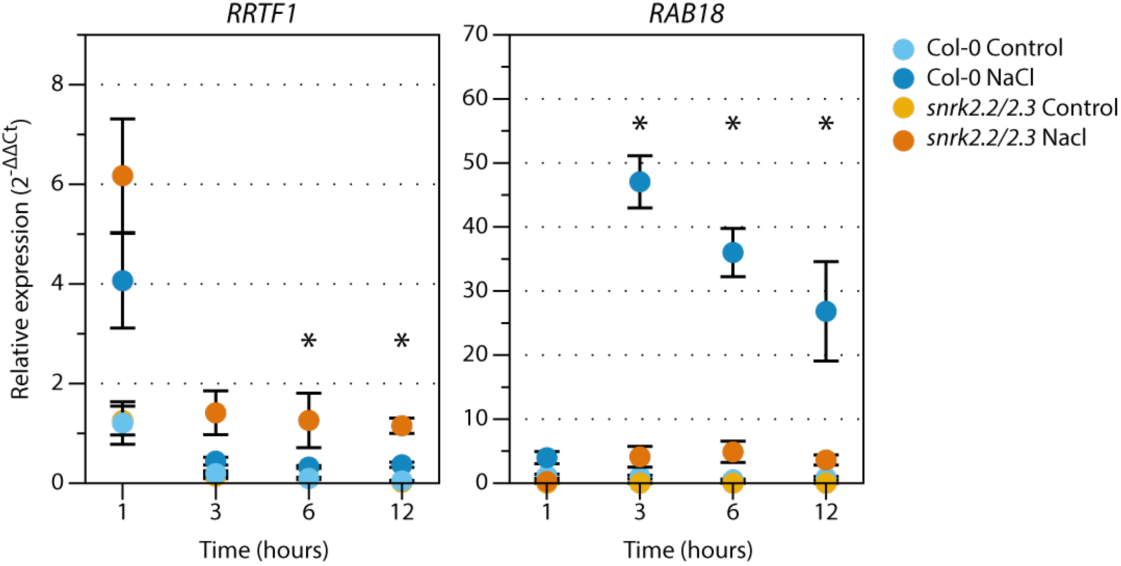
ABA signaling mutants show enhanced *RRTF1* expression at 3 hours after salt treatment onwards. (A) *RRTF1* and *RAB18* expression in 7-day-old *snrk2.2/2.3* and wild type seedlings, transferred to 125 mM NaCl or control (1/2 MS) medium, and harvested after multiple timepoints. Levels are normalized for control conditions at 1 hour. Data was statistically analyzed using two-way ANOVA per timepoint, followed by a Tukey post-hoc test. Letters of the same font indicate significance (p < 0.05).

**Figure S10.**
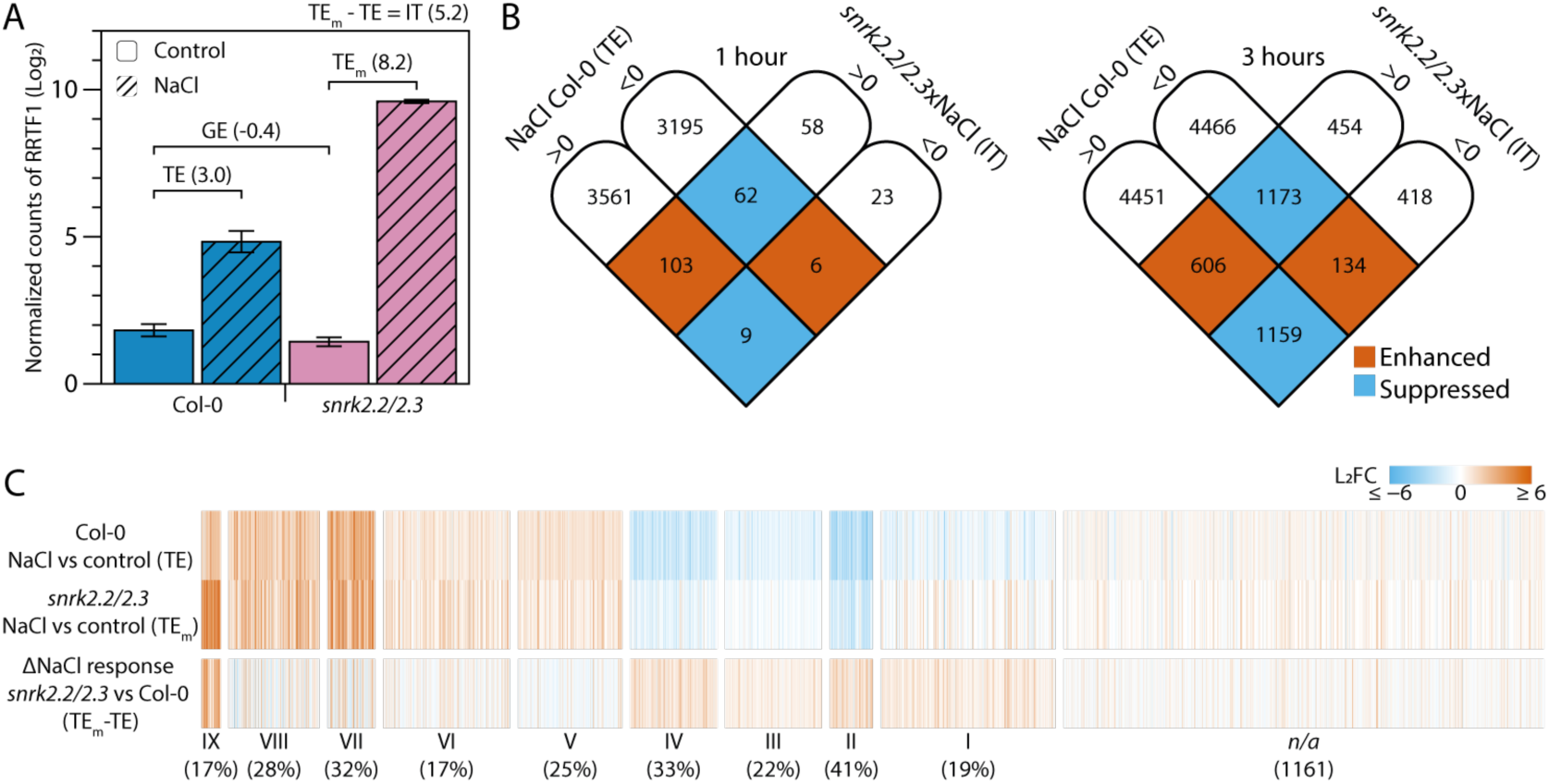
The ABA-insensitive *snrk2.2/2.3* mutant shows enhanced sodium-induced transcriptional responses. (A) Example of normalized counts of *RRTF1* in *snrk2.2/2.3* and wild type Col-0 at 3 hours. Different outputs of the DESeq2 software are indicated. Treatment effect (TE; response to a treatment in wildtype), Genotype effect (GE; difference between mutants and wildtype in control conditions) or Interaction term (IT; differences in response to a treatment between a mutant and wildtype). The IT is the difference between the treatment effect of the mutant (TE_m_) and the TE. Bars represent averages +/- SE (n=3). (B) The overlap between the *snrk2.2/2.3*xNaCl IT with NaCl and the NaCl TE (Col-0) for up- and downregulated DEGs (FDR < 0.05) at 1 and 3 hours after stress induction. Colors indicate enhancement or suppression. (C) The full heatmap of Fig. 5B including the genes that were identified in the RNA sequencing experiment of Fig. 5, but not in the experiment of Fig. 1.

**Figure S11.**
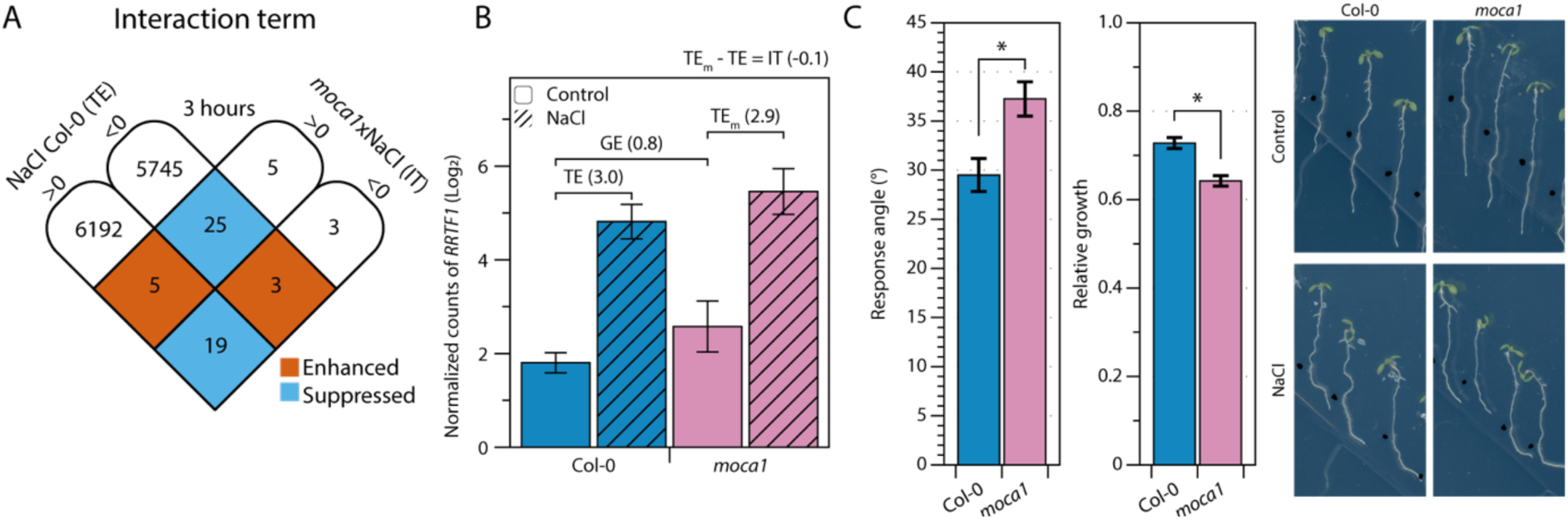
The *moca1* mutant is not impaired in halotropism, and shows very few differential transcriptional responses during NaCl stress. (A) The overlap between the *moca1*xNaCl IT with NaCl and the NaCl TE (Col-0) for up- and downregulated DEGs (FDR < 0.05) at 3 hours after stress induction. Colors indicate enhancement or suppression. (B) Normalized counts of *RRTF1* in Col-0 and *moca1*. Bars represent averages +/- SE (n=3). Different outputs of the DESeq2 software are indicated. Treatment effect (TE; response to a treatment in wildtype), Genotype effect (GE; difference between mutants and wildtype in control conditions) or Interaction term (IT; differences in response to a treatment between a mutant and wildtype). The IT is the difference between the treatment effect of the mutant (TE_m_) and the TE. Bars represent averages +/- SE (n=3). (C) Quantification and representative images of the halotropism response and relative root growth (NaCl/control) of 7-day old Col-0 and *moca1* seedlings at 48 hours after the introduction of the NaCl gradient. Data points represent averages +/- SE, n=40-48. Letters indicate significant differences after Dunn test (p < 0.05). Brackets and asterisks indicate significant differences using unpaired T-tests for unequal variance (* = p < 0.05).

**Figure S12.**
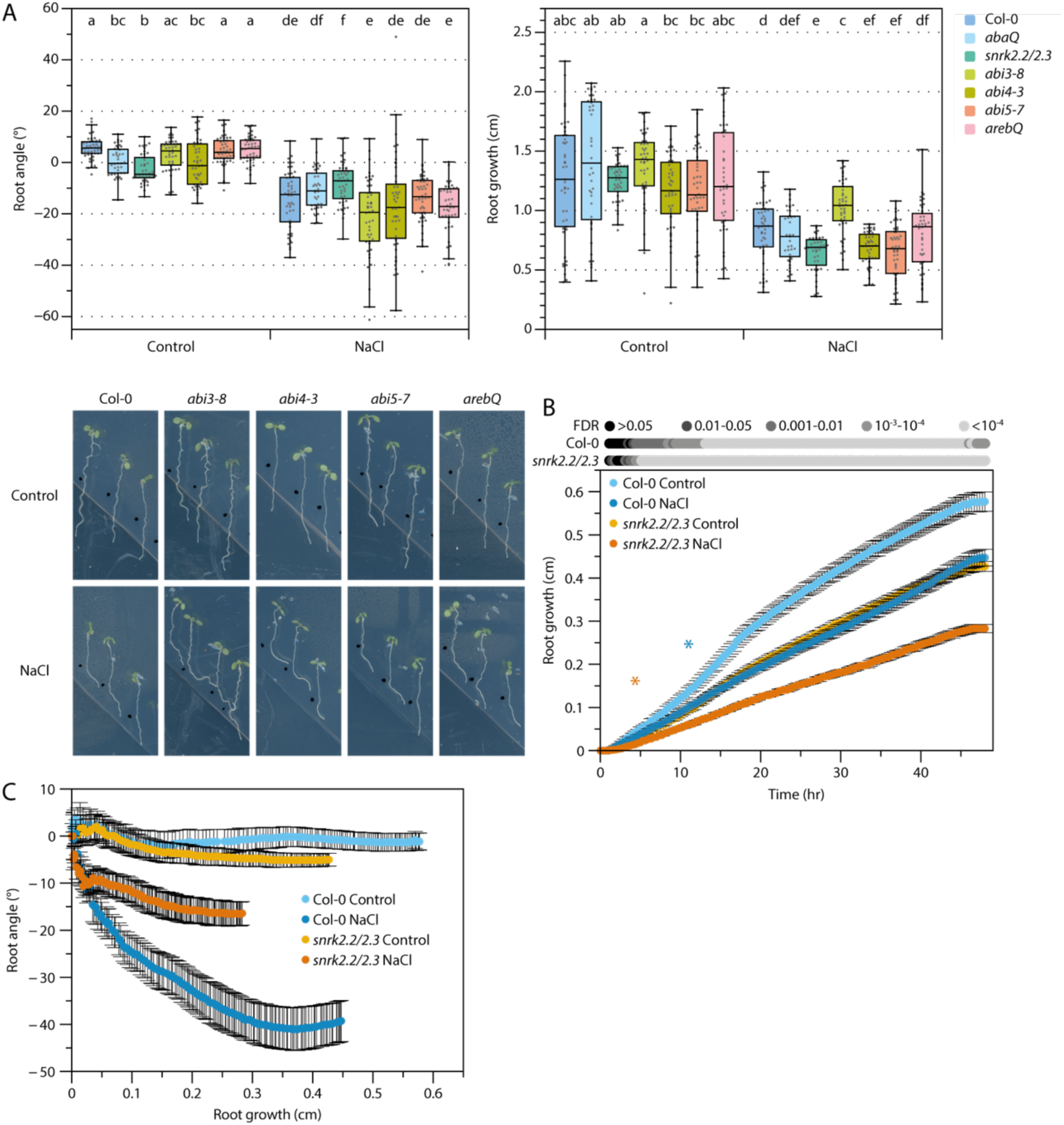
ABA signaling mutants show a reduced halotropic response. (A) Absolute data and representative images of the halotropism assay shown in Fig. 5A. Statistical analysis was performed using non-parametrical Dunn’s test and corrected for multiple testing using Benjamin-Hochberg (BH) procedures. Letters indicate statistical groups. (B-C) Absolute data and representative images of the halotropism assay shown in Fig. 6B. Seedlings were imaged every 20 minutes for 48 hours. Dots represent mean values (n=20) +/- SE. Pairwise Welch tests were performed for NaCl treated samples and the control condition of the same genotype and corrected for multiple testing with BH. Corrected p-values (NaCl vs control) are shown in grey-values above the graph. (C) Timelapse data as shown in Fig. 5B, plotted with the angle against the length.

**Figure S13.**
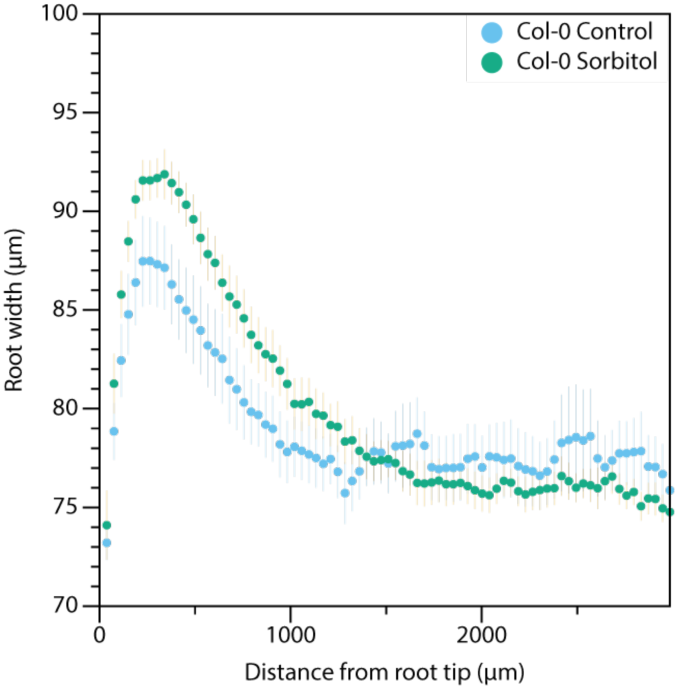
Root width is not increased by osmotic stress. Root width along the root axis (length 0 = root tip) for Col-0 with control or 250mM Sorbitol treatment. No significant differences were observed using multiple Welch tests followed by BH correction for multiple testing (Sorbitol vs control). (n = 10).

**Figure S14.**
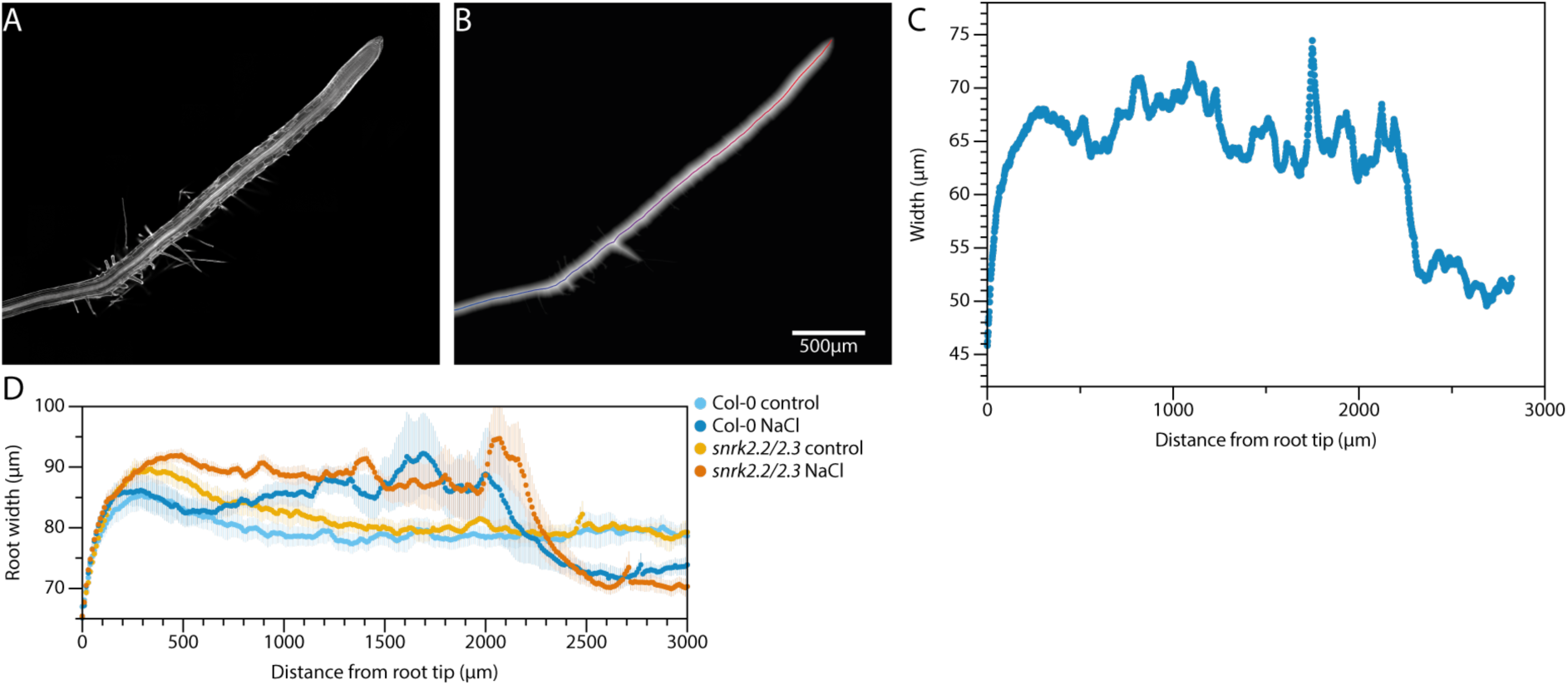
Root width was analyzed using an automatic pipeline. (A) Confocal image of a NaCl-treated root fixed at 24 hours after stress induction and counterstained with Calcofluor white. (B) Euclidean distance map of a segmented image. Roots were segmented based on pixel values, followed by calculating the distance of each root-pixel to the closest non-root pixel (indicated by gray intensity). The midline was calculated using this distance map, filtering for the highest distance-values in an area of 5×5 and connecting these filtered pixels together. The midline is indicated from red to blue (red = root tip). Next, the Euclidean distance values of every point on the midline were used as root width. (C) Plot of the root width along the midline. (D) Quantification of root width ratio along the root axis (length 0 = root tip) for Col-0 and *snrk2.2/2.3*. The region of 0-1400 µm is shown in Fig. 6D. NaCl induced cell swelling from the root tip till 2200 µm from the root tip. Older cells of the root were shrunken by the salt treatment (2200-3000 µm).

**Figure S15.**
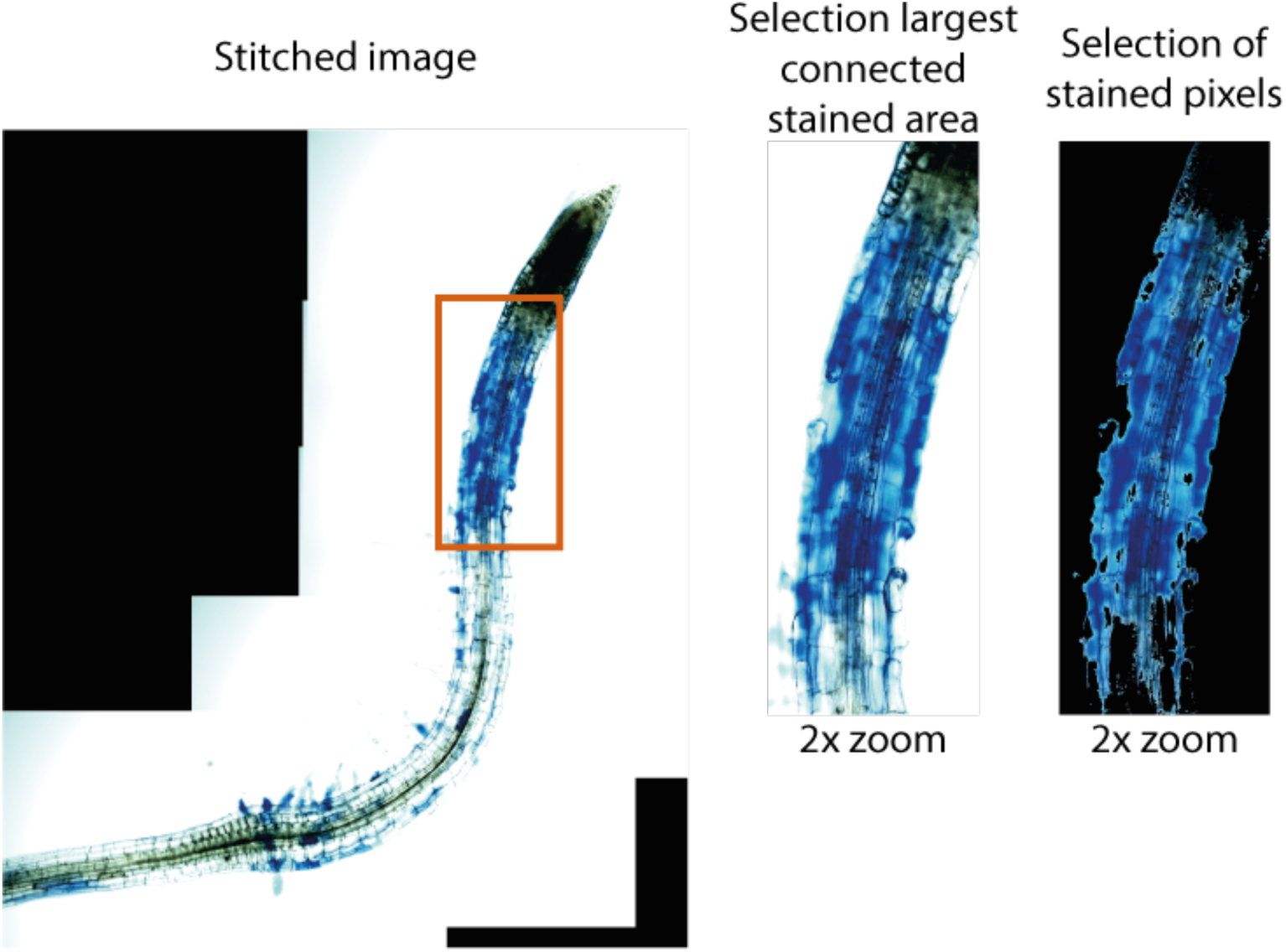
Evans Blue staining was analyzed with a script. Automized script to analyze the Evans Blue staining. First pixels are filtered to be dominantly blue (in an RGB channel), followed by the selection of the largest continuous blue object and all object in close vicinity (displayed in the 2^nd^ panel). Finally, all blue pixels in this selected area are counted (displayed in the 3^rd^ panel).

